# Characterization of Cell-cell Communication in Autistic Brains with Single Cell Transcriptomes

**DOI:** 10.1101/2021.10.15.464577

**Authors:** Maider Astorkia, Herbert M. Lachman, Deyou Zheng

## Abstract

Autism spectrum disorder is a neurodevelopmental disorder, affecting 1-2% of children. Studies have revealed genetic and cellular abnormalities in the brains of affected individuals, leading to both regional and distal cell communication deficits. Recent application of single cell technologies, especially single cell transcriptomics, has significantly expanded our understanding of brain cell heterogeneity and further demonstrated that multiple cell types and brain layers or regions are perturbed in autism. The underlying high-dimensional single cell data provides opportunities for multi-level computational analysis that collectively can better deconvolute the molecular and cellular events altered in autism. Here, we apply advanced computation and pattern recognition approaches on single cell RNA-seq data to infer and compare inter-cell-type signaling communications in autism brains and controls. Our results indicate that at a global level there are cell-cell communication differences in autism in comparison to controls, largely involving neurons as both signaling senders and receivers, but glia also contribute to the communication disruption. Although the magnitude of change is moderate, we find that excitatory and inhibitor neurons are involved in multiple intercellular signaling that exhibit increased strengths in autism, such as NRXN and CNTN signaling. Not all genes in the intercellular signaling pathways are differentially expressed, but genes in the pathways are enriched for axon guidance, synapse organization, neuron migration, and other critical cellular functions. Furthermore, those genes are highly connected to and enriched for genes previously associated with autism risks. Overall, our proof-of-principle computational study using single cell data uncovers key intercellular signaling pathways that are potentially disrupted in the autism brains, suggesting that more studies examining cross-cell type affects can be valuable for understanding autism pathogenesis.

## Introduction

Autism spectrum disorder (ASD) is a class of neurodevelopmental disorders characterized by two main clinical and behavior domains: i) social communication and interaction deficits and ii) restricted interests and repetitive behaviors, according to DSM-5 classification^1^. The Centers for Disease Control and Prevention (CDC) estimate that ASD prevalence is 1 in 54 in children aged 8 years old and ∼4 times more in boys than in girls, based on data from a 2016 report^2^. Genetics is the major risk factor with contributions from both rare and common, inherited and *de novo* variants, but environment also plays significant roles. To date, the Simons Foundation Autism Research Initiative (SFARI)^3^ has collected 1,003 genes related to ASD risk supported by various levels of evidence, but individually each of the genes accounts for less than one percent of the cases. These genes have a diverse array of functions but can be categorized into several broad molecular and cellular pathways, including synaptic function, signaling pathways (such as WNT, bone morpho-genetic protein (BMP), Sonic hedgehog (SHH) and retinoic acid (RA)), and chromatin remodeling^4–7^. Overall genetic studies suggest that the neuropathological mechanisms behind ASD are complex and multifaceted, and the risk genes have roles in multiple molecular and cellular networks.

Mutations in ASD risk genes can lead to structural and functional disruptions in neural circuits. At the genetic and cellular levels, synaptogenesis and synapse function have been a long and major focus. Cell-cell signaling regulate synapse formation and plasticity but they are also key for other cellular processes such as immune response, cell differentiation and maturation, and cell homeostasis^8,9^. Many studies have pointed to mutations in ASD risk genes related to such intercellular communications. Carias and Wevrick^10^ studied 94 genes with *de novo missense* (DNMs) variants in Tourette Syndrome and ASD and found that many of them are membrane-associated proteins, cell-cell adhesion or communication proteins, G-protein signaling, and centrioles or cilia regulators or components. Similarly, analyzing data from Whole Exome Sequencing (WES) and Whole Genome Sequencing (WGS), Zhang *et al*. found that many DNM ASD candidate genes are involved in cell-cell communications ^11^. Furthermore, in a Genome-Wide Association Study (GWAS) involving 18,381 ASD and 27,969 control individuals, Grove *et al*.^12^ identified five genome-wide significant loci and additional strong candidates, including genes involved in cell interaction and signaling (*NEGR1, GADPS, and KCNN2*). A subsequent WES study reporting 102 risk genes also supported the roles of ASD genes in gene regulation and neuronal communication, and further linked some genes to delayed age of walking and reduced IQ^13^. In a separated study focused on ultra-rare variants, Wilfert *et al*.^14^ also uncovered risk genes implicated in intercellular signaling (e.g., Erb signaling). Non-genetic studies have also pointed to structural changes in ASD brain that can lead to disrupted circuits and connections. For example, in 2009 Keary *et al*.^15^ and Casanova *et al*.^16^ showed that corpus callosum volume in ASD brains were reduced and gyral window was abnormally narrow. A recent neuroimaging and computational study from the Autism Brain Imaging Data Exchange Initiative further revealed both microcircuit and macroscale connectome abnormality in autism brains^17^. Chien *et al*. also observed thinner cortical thickness in bilateral cingulate subregions in ASD individuals^18^.

To investigate cell communication appropriately, studies need to be performed at individual cell or cell type level. This is particularly important for brain disorders because a human brain is composed of many types and subtypes of neurons and glia that are located in multiple brain layers and regions, with some exhibiting layer and regional specificity. Several studies have investigated cell-type and brain-layer expression patterns of risk genes for ASD and other psychiatric disorders. For example, we previously analyzed multiple *in vitro* and *in vivo* human neural and brain expression datasets from both control and ASD studies and demonstrated that inhibitory neurons could be the major cells affected in ASD, supporting an imbalance between inhibitory and excitatory signaling as important for ASD pathogenesis ^19^. DNM variants disrupting protein-protein interactions were identified in excitatory and inhibitory neuronal lineages by Chen *et al*.^20^. Likewise, in a WES study, Satterstrom *et al*.^13^ reported enriched expression of the 106 ASD risk genes in maturing and mature excitatory and inhibitory neurons, as well in oligodendrocyte progenitor cells and astrocytes. In addition, studies from the PsychENCODE Consortium further underscored cell specific roles^21,22^. Most recently, Velmeshev *et al*.^23^ applied single cell transcriptomics to compare ASD brains to controls and showed that gene expression changes occurred predominantly in microglia and upper layer excitatory neurons, while affected genes were enriched for synaptic function. Furthermore, they suggested that ASD severity was significantly linked to groups of genes expressed in specific neurons. Reanalyzing the same data, Ji *et al*. identified cell-type-specific ASD-associated gene modules and showed that excitatory neurons, such as those of layer 2/3 (L2/3), L4 and L5/6-CC (cortico-cortical projection neurons) could play essential roles in ASD^24^.

While these studies have underscored the roles of specific cell types, a direct and detailed examination of cell-cell communications in ASD brains has not been reported. On the other hand, cell-cell communication (CCC) has become a new computational frontier in single cell transcriptomic (scRNA-seq) studies, with the goal to systematically characterize intercellular communications between cell types^25–27^. This kind of studies are typically focused on molecular interactions involved in ligand-receptor, receptor-receptor and extracellular matrix-receptor proteins. The interactions can be categorized into four main types: 1) Autocrine signaling for intracellular communication, 2) paracrine communication in which molecules are secreted by diffusion, 3) juxtracrine communication based on direct contact established by gap junctions, and 4) endocrine where molecules are secreted and signaling is achieved through extracellular fluids^9,28^. Since these communications rely on the organization of cellular activities, it is important to consider the expressed molecules, associated pathways, the directionality, the magnitude, and the biological relevance^9^. Many software have been developed to use single cell transcriptomic data for predicting cell-cell communications, mostly based on cell type (or subtype) annotation and a curated database of ligand-receptor and other interacting surface proteins^29–32^. The databases and the computational methods for inferring CCC, however, are quite diverse.

After evaluating multiple software packages and considering their benchmark performances^26,33^, we chose CellChat^32^ for inferring, visualizing and comparing CCC in this study, mostly because it provides robust tools for multiple level of CCC analyses. Its database consists of 2,021 validated and curated protein interactions, including paracrine/autocrine ligands/receptors, extracellular matrix receptor interactions, and cell-cell contact interactions. In contrast to other software, it has an advantage in taking into account the heterodimeric complexes involved in cell-cell interactions, as well as soluble agonists, antagonists, and stimulatory and inhibitory membrane-bound co-receptors^32^. Additionally, CCC is calculated using mass action model and its prediction can be used for network analysis, patter recognition methods and manifold learning approaches. More specifically, we applied CellChat to a previously published single cell nuclei RNA-seq (snRNA-seq) data from 15 autistic and 16 control brains. In the voriginal study, Velmeshev *et al*. have analyzed differentially expressed genes (DEGs) in the 17 cell types and their enriched functions, suggesting that upper-layer excitatory neurons and microglia are preferentially affected in autism^23^. While the authors identified dysregulated genes involved in synaptic function, such as *SYN1* and *NRXN1*, a systematic analysis of cell-cell interactions was not explored. Here, we conducted a comparison of the cell-cell communication networks in the two groups of brains, and then identified critical intercellular signaling that were altered in the ASD brains. In addition, we analyzed function enrichment of the genes within the altered signaling pathways, their relationship to known ASD risk genes, and their connection to differential intracellular pathways in ASD brains compared with controls.

## Materials and methods

### Human brain single nucleus RNA-seq data

The human snRNA-seq data, consisting of 104,559 nuclei from prefrontal (PFC) and anterior cingulate cortex (ACC) post-mortem tissues^23^, were downloaded from the UCSC cell browser (https://cells.ucsc.edu/?ds=autism). The gene expression matrix was already normalized by total unique molecular identifiers (UMIs) per nucleus and log2-transformed. PFC data consisted of 62,166 cells from 13 ASD (32,019 cells) and 10 control samples (30,147 cells), while 9 ASD (19,984 cells) and 3 Control (22,409 cells) ACC samples made up to 42,393 nuclei. We directly used the authors’ original nuclei classification of 11 neuronal types: parvalbumin interneurons (IN-PV), somatostatin interneurons (IN-SST), SV2C interneurons (IN-SV2C), VIP interneurons (IN-VIP), layer 2/3 excitatory neurons (L2/3), layer four excitatory neurons (L4), layer 5/6 corticofugal projection neurons (L5/6), layer 5/6 cortico-cortical projection neurons (L5/6-CC), maturing neurons (Neu-mat), NRGN-expressing neurons (Neu-NRGN-I), NRGN-expressing neurons (Neu-NRGN-II), and 6 non-neuronal cell types: fibrous astrocytes (AST-FB), protoplasmic astrocytes (AST-PP), oligodendrocyte precursor cells (OPC), oligodendrocytes, microglia cells and endothelial cells.

### Cell-cell communication analysis in PFC data

We performed cell-cell communication analysis using the CellChat^32^ software, with PFC and ACC samples separately. For every pair of cell types, ligand-receptor (L-R; and other) interactions were identified and measured. Intercellular communication is based on the projected ligand and receptor profiles where the expression level of L and R is approximated by their geometrical mean across individual cells of a type. These interactions represent the interaction strengths (also referred as “probability” in CellChat) between all ligand-receptors expressed in two given cell types. Note that CellChat considers important signaling factors such as heteromeric complexes involved in each interaction in addition to a L-R pair, therefore, the absence of any of those components leads to a null interaction. Genes expressed in less than 20% of cells in one cell type were excluded and only statistically significant (p < 0.05, permutation test) communications were considered in our analysis.

We took two complementary approaches to define cell-cell interactions in ASD and controls and differences between them (**Figure 1a**). In the “Global approach”, cells from all ASD samples were merged into one group and the control cells into another; ligand-receptor interactions were calculated and compared between the two groups. From individual L-R interactions to signaling, interaction score of a signaling pathway was calculated by summing up the interaction strengths for all the L-R in the pathway. Dysregulated signaling pathways between ASD and controls were then identified by CellChat (Wilcoxon test, p < 0.05). CellChat also allows the identification of key signaling and latent communication patterns across all signaling pathways. It applies Cophenetic and Silhouette metrics, both based on hierarchical clustering, to identify the numbers of patterns as well as major sending/outgoing and receiving/incoming signaling pathways for both conditions. In the second “Sample-by-sample approach”, we treated each sample independently, i.e., the strengths of L-R interactions and signaling pathway scores were calculated for each sample. After pathways identified in less than 5 samples were removed, significantly dysregulated L-R interactions in the remaining pathways for each pair of cell types were identified by applying Wilcoxon test (p < 0.05) to the sample level data (i.e., “pseudo-bulk values”), comparing ASD to controls. Among the significantly differential pathways, we picked 7 with literature support for their involvement in ASD, and then used chord diagrams to illustrate the cell-cell communications and differences between ASD and controls. Chord diagrams were based on the differences in ASD vs control pathway strengths between individual pairs of cell types.

**Figure 1.**
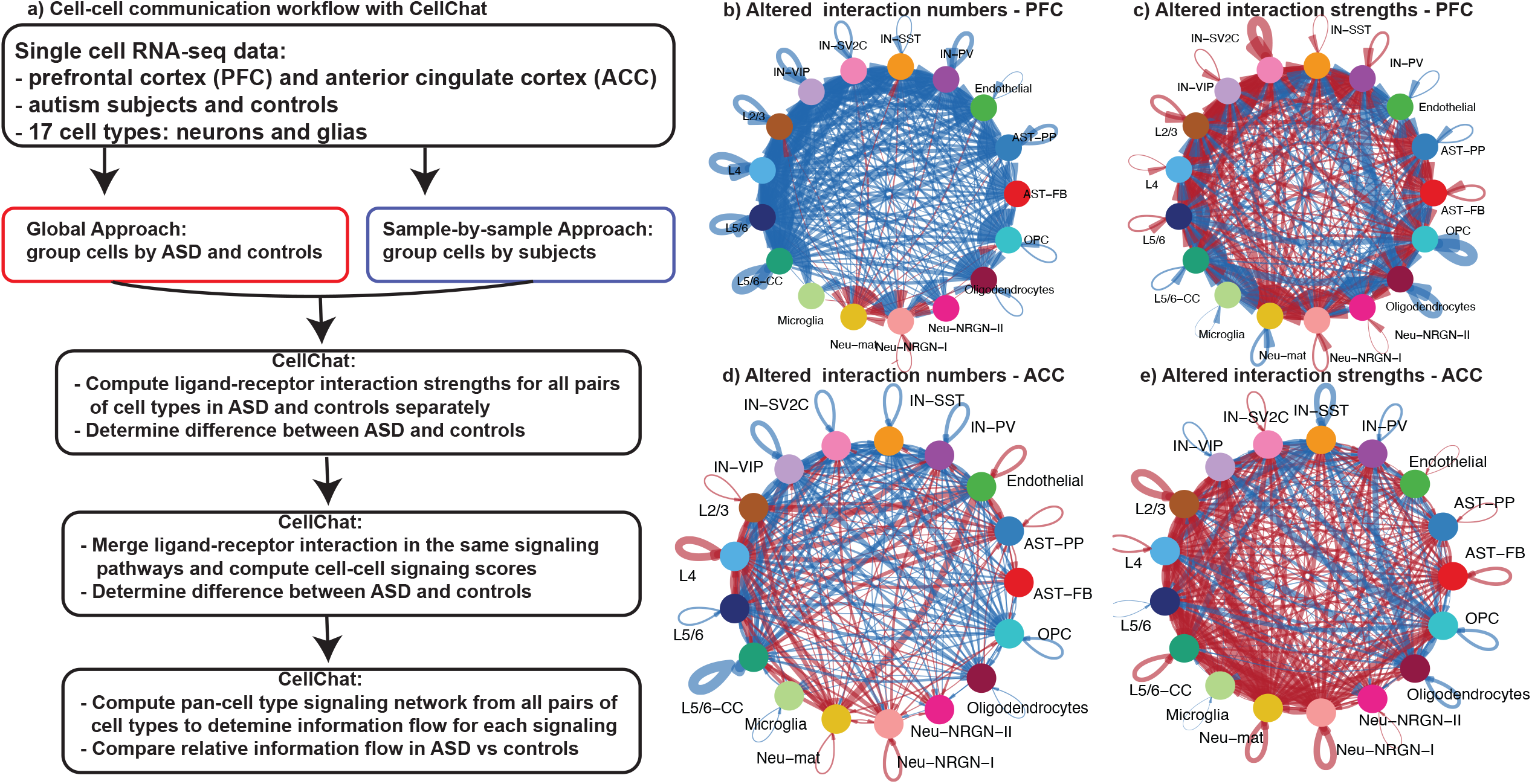
Change in communication between individual pairs of cell types in ASD. a) Computational workflow of key analytic steps. b) Difference in the total numbers of L-R interactions in ASD vs control PFC. c) Difference in the total strengths of L-R interactions in ASD vs control PFC. d) Difference in the total numbers of L-R interactions in ASD vs control ACC. e) Difference in the total strengths of L-R interactions in ASD vs control PFC. In b-e, lines indicate the changes in individual pairs of cell types, with red for increase and blue for decrease in ASD. The thickness of the lines represents the extent of changes, with the maximal corresponding to 28 in b/d and 0.36 in c/e.

### Over representation analysis based on curated gene sets and Gene Ontology

Two over representation tests were performed to test statistical enrichment for genes either in selected Gene Ontology (GO) terms or in previously curated gene sets related to brain disorders, using all genes involved in the disrupted pathways as the test set. In the former, cell-cell interacting genes in dysregulated pathways were used for enrichment of GO “Biological Processes” terms (false discovery rate (FDR) < 0.05). EnrichGO function from clusterProfiler^34^ was used to run this and “simplify function” was used to reduce redundancy in enriched GO terms. We also applied ClueGO (v2.5.8)^35^ to the most significantly enriched GO terms (FDR < 0.0005) to better define function enrichment and to reduce redundancy. In the latter, we first obtained various gene sets. ASD candidate genes were downloaded from two sources: 1) SFARI database (https://www.sfari.org) (ASD_SFARI) comprising 1,003 genes scored as syndromic, high confidence, strong candidate and suggestive evidence levels, and 2) AutismKB^36^ (ASD_AutismKB) with 228 genes. Two schizophrenia (SCZ) risk gene sets were obtained from SZDB2.0 (http://www.szdb.org/index.html) database; SCZ_GWAS gene set was built with 571 genes identified in GWAS studies and SCZ_CNV gene set with 408 genes affected by Copy Number Variation (CNV). The 1190 genes related to bipolar disorder were obtained from a previous study^19^, while gene sets for intellectual Disability (ID_CNV; n=908) and Attention Deficit Hyperactivity Disorder (ADHD; n=359) were obtained from (http://www.ccgenomics.cn/IDGenetics/index.php) and (http://adhd.psych.ac.cn/index.do) databases, respectively. These 7 gene lists were then used for over representation analysis of genes in dysregulated pathways using the Fisher’s test in the GeneOverlap package^37^ in R.

### Protein-protein interaction network

The STRING database^38^ was used to identify the protein-protein interaction (PPI) networks connecting the SFARI ASD genes to the genes in the dysregulated CCC signaling pathways. The sources to use for PPI were experiments, databases, co-expression, gene fusion and co-occurrence databases. A highest confidence of 0.9 was applied and disconnected nodes were removed.

### Gene enrichment analysis based on cell types

To define pathways significantly affected by expression changes between ASD and controls, for each cell type we applied Gene Set Enrichment Analysis (GSEA, v4.1.0)^39,40^ to all genes expressed in that cell type, ranked by their expression fold changes between ASD and control samples. The enriched GSEA sets (FDR < 25%) in the GO “Biological Processes” were retrieved to determine the significant GO terms containing genes encoding ligands or receptors.

### Spatial co-expression of ligands and receptors

Maynard *et al*.^41^ generated spatial gene expression profiles for six layers and white matter of two pairs of “spatial replicates” from three dorsolateral prefrontal cortex samples^41^. We used these data to compute Pearson’s correlation coefficient between a pair of L-R across cells in all layers or within specific layers, with the results compared to those from randomly created L-R pairs.

### Cell-cell communication analysis in Anterior Cingulate Cortex data

We applied the above analysis for the PFC samples to the ACC data, with the same designs and same parameters.

## Results

### Numbers and strengths of L-R interactions in autistic PFC and controls

Most current computational studies of cell-cell communications (or interactions) from single cell transcriptomics data analyze ligand-receptor interactions among all cell types in one sample, but some software also provides methods for comparing interactions between two samples. We thus started with this common practice by comparing cell-cell interactions between ASD and control brains using a previously published scRNA-seq data^23^ (**Figure 1a**). The data contained samples from prefrontal (PFC) and anterior cingulate cortex (ACC), while cells/nuclei were classified into 17 types (see Methods). We will focus our description on PFC below as its relation to social and cognition function and ASD is relatively better studied, but the same bioinformatics methods were applied to ACC data. We merged cells from all 13 ASD PFC samples into one group and cells from 10 control PFCs into the other, and then studied the between-group difference (referred as “Global approach”; **Figure 1a**). We used the software CellChat, which analyzes the expression of ligands, receptors and other proteins involved in cell-cell interactions to determine the interaction strength in an interacting protein complex for a pair of cell types^32^. For simplicity, we hereafter refer all pairwise interactions as ligand-receptor (L-R) interactions even though non-canonical ligands or receptors are also considered. In total, CellChat identified 6,433 and 5,049 interactions in control and ASD PFC, respectively. The highest number of interactions in controls was 103, found in the L5/6-CC to L5/6-CC (cortico-cortical projection neurons) autocrine interactions, while the highest in ASD sample was 79, between L5/6-CC and L2/3 cell types. The total interactions in ACC are similar to those in PFC, with 5,940 in control and 5,843 in ASD. In both control and ASD ACC, L5/6-CC autocrine signaling had the most interactions, 126 in control and 109 in ASD. In both tissues, L5/6-CC and L5/6 (layer 5/6 corticofugal projection neurons) are the cell types with the most interactions as either sender or receiver cells (maximal of 904 and 771 in L5/6-CC and L5/6, respectively), while microglia and Neu-NRGN-II have the least (maximal of 37 and 96 in microglia and Neu-NRGN-II, respectively). This difference is not directly related to cell numbers, because there is no significant correlation between cell numbers and L-R interaction numbers (Pearson’s correlation coefficient = 0.07) (**Figure S1**).

Although moderate (largest reduction is 28, seen between L4 and L5/6 CC), we found a decreased trend in the numbers of cell-cell interactions in ASD vs controls for the 17 cell types in PFC (**Figure 1b; Figure S2a**), involving both autocrine and paracrine signaling. Although this analysis indicates reduced interactions in ASD for most cell types except two (Neu-mat and Neu-NRGN-I), it did not account for the interaction strengths, which capture the actual expression levels of all the genes encoding proteins in an interaction. With strengths of all L-R interactions considered (the maximal values are 0.37 in control and 0.4 in ASD), we found that cell-cell communications between many pairs of cell types became overall stronger in ASD (156 increased vs 133 decreased), especially when excitatory and inhibitory neurons were involved (**Figure 1c; Figure S2b**). At the level of individual L-Rs, the interaction strengths also exhibited a trend of moderately increased in ASD (data not shown). The largest increase was found in an autocrine signaling for IN-SV2C interneurons, followed by signaling between IN-SV2C inhibitory neurons and L2/3 and L4 excitatory neurons, suggesting potential abnormal excitatory-inhibitory neuronal communications in ASD brains. Among the decreased interactions in ASD, paracrine signaling between oligodendrocyte and two inhibitory neurons (IN-SV2C and IN-PV) showed the most reduction, followed by OPC autocrine signaling. These findings are very interesting, not only in terms of major implications of inhibitory neurons but also with respect to our increasing appreciation of the role of glia in neuron development and functions, such as synapse modification^42,43^. In addition, reduction of oligodendrocyte cells was previously reported and linked to neurodevelopmental disorders^44,45^.

The ACC shows a different pattern of alterations in cell-cell interactions, with similar numbers of cell type pairs exhibiting increased or decreased interactions in ASD brains (**Figure 1d,e; also see below**).

### Major intercellular signaling patterns in PFC

Next, we grouped L-R interactions into signaling pathways using CellChat and compared them between ASD and controls. This is a key advantage of CellChat over other software, and it is important because many L-R interactions would invoke the same downstream signaling, such as FGF1/2/3-FGFR1/2/3 or NRNX1/2/3-NLGN signaling. Afterwards, we used CellChat to identify the major latent patterns in outgoing and incoming signaling in the control and ASD PFC, by clustering cell types sending or receiving similar signaling. For outgoing signaling (**Figure 2a,b**), CellChat uncovered 5 and 4 patterns for control and ASD samples, respectively. In control (**Figure 2a**), pattern 1 involved interneurons, Neu-mat and Neu-NRGN-I cells, in which the main pathways were IGF, TAC, HGF and OSTN. Pattern 2 involved solely excitatory layer neurons with NOTCH, WNT, EPHB and NT as main contributors. Oligodendrocytes clustered by themselves in Pattern 3, expressing SPP1, CD22 and MAG. Pattern 4 involved a mixture of non-neuronal and neuronal cells with endothelial, microglia. and Neu-NRGN-II cells, with FN1, ESAM, OCLN and PECAM1 as the coordinators. Finally, pattern 5 was comprised of only non-neuronal astrocytes and OPC cells, in which the expression of genes related to VEGF and ANGPTL were the most abundant. On the other hand, pattern 1 from ASD PFC was a mixture of non-neuronal and neuronal cells with astrocytes, interneurons, and Neu-mat (**Figure 2b**). In this pattern, genes encoding IGF, CRH, BMP and TAC signaling were the dominant ligands. In pattern 4, similarly, both non-neuronal and neuronal cells were involved, with a mixture of microglia and NRGN-expressing neurons. Here, CD39, CD45 and VISTA were the main senders. Like controls, excitatory neurons grouped together in pattern 2 with outgoing signals from NT, ANGPT and CSF pathways. Pattern 3 clustered only non-neuronal cell types, endothelial and oligodendrocytes cells, with CD22, MAG, CLDN and ESAM as the main signaling contributors.

**Figure 2.**
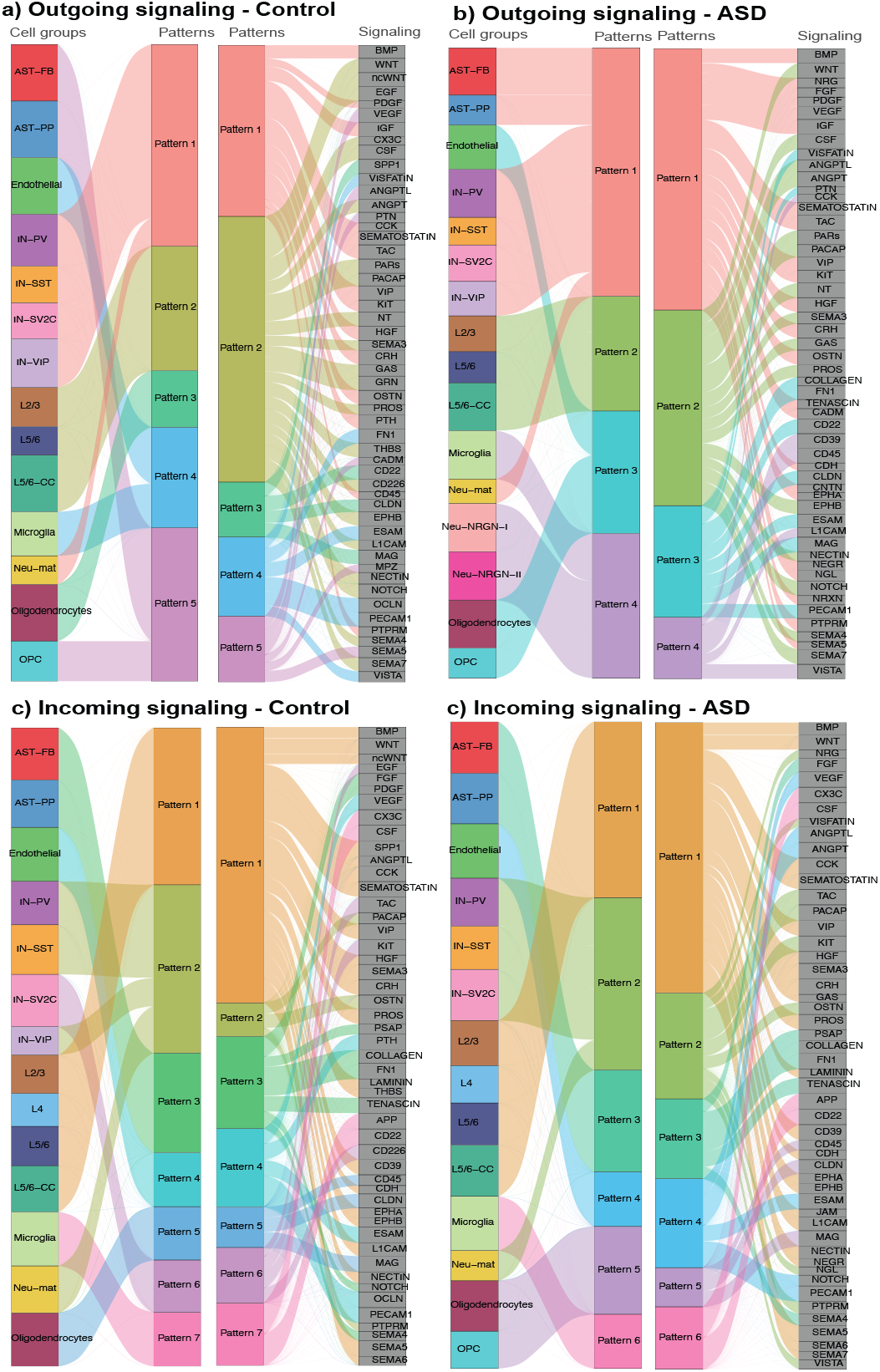
Cell-cell communication patterns in PFC. The networks show the CellChat inferred latent patterns connecting cell groups sharing similar signaling pathways. The thickness of the water flow represents the relative contribution of the cell group or signaling pathway to a latent pattern; outgoing patterns for secreting cells in control (a) and ASD (b); incoming patterns of receiving cells in control (c) and ASD (d).

From the receiving end, 7 patterns were found in control PFC, while 6 in ASD (**Figure 2c,d**). In control (**Figure 2c**), pattern 1 involved excitatory layer neurons and Neu-NRGN-I neurons and others, while SEMA5, VIP and WNT were the main signaling receivers. Pattern 2 grouped interneurons and Neu-mat cells with genes related to OSTN and PACAP pathways. FN1 and TENASCIN pathways were the main signal receivers in pattern 3, in which astrocytes and OPC participated. Endothelia, oligodendrocytes and microglia clustered by themselves in patterns 4, 5 and 7. Finally, IN-SV2C and Neu-NRGN-II cell types grouped in pattern 6 with TAC, KIT and CD226 as main receptors for incoming signals. For ASD (**Figure 2d**), pattern 1 was formed by excitatory layer neurons and Neu-NRGN-I cells with SEMA5, CCK and SEMATOSTIN pathways as the main receptors. Interneurons and Neu-mat cells clustered again together in pattern 2, with TAC, KIT and PACAP pathways. Genes involved in FN1 and TENASCIN pathways were the predominant receptors in pattern 3, enclosing astrocytes and Neu-NRGN-II. Endothelial, both oligodendrocyte cell types and microglia were enclosed in pattern 4, 5 and 6, respectively, and showed same patterns as in control dataset.

Taken together, analysis of the intercellular CCC patterns indicates that all cell types contribute, although at various extents, in both control and ASD PFCs, and no cell type appears to serve as major signaling hubs in either condition. The CCC patterns seem similar in ASD and controls, but it appears a little less diverse (one fewer pattern) in ASD.

### Pan-cell type signaling difference between ASD and control PFCs

Considering that one ligand may interact with receptors on multiple cell types, and vice versa, we next examined intercellular signaling for which multiple or all 17 cell types participated, in contrast to looking at cell-cell communications between any two cell types as described above. Thus, we used CellChat to sum up all the L-R interaction strengths in the same signaling pathway across all cell types in which the L-R encoding genes were expressed. In total, CellChat has a collection of 229 families of signaling pathways. Among them, 68 unique ones were identified in our global approach. CellChat considers this pan-cell CCC network as “information flow.” Applying it to our data, we obtained “relative information flow” change between ASD and controls (**Figure 3a**). The result indicates that 8 signaling pathways, non-canonical WNT (ncWNT), GRN, EGF, THBS, PTH, OCLN, SPP1 and CD226, were specifically identified in the control dataset but no pathways unique to ASD were found. Additionally, 48 of the signaling pathways were significantly down-regulated and 6 up-regulated in ASD when the relative information flow was analyzed statistically (**Figure 3a**; p < 0.05, Wilcoxon test). However, we should mention that the differences for most signaling are relatively small. Close examination of the changes in relative contributions of individual cell types with respect to both outgoing and incoming signaling in ASD and control datasets illustrates this better (**Figure 3b,c)**. It also indicates that the two largest reduced signaling pathways (ncWNT and THBS) are due to significant decreases in incoming signaling in multiple cell types. This close-in analysis further suggests that excitatory and inhibitory neurons are the major sources of signaling senders while both neurons and glial cells participate as receivers. The contribution of microglia to information flow, however, appears low in this analysis.

**Figure 3.**
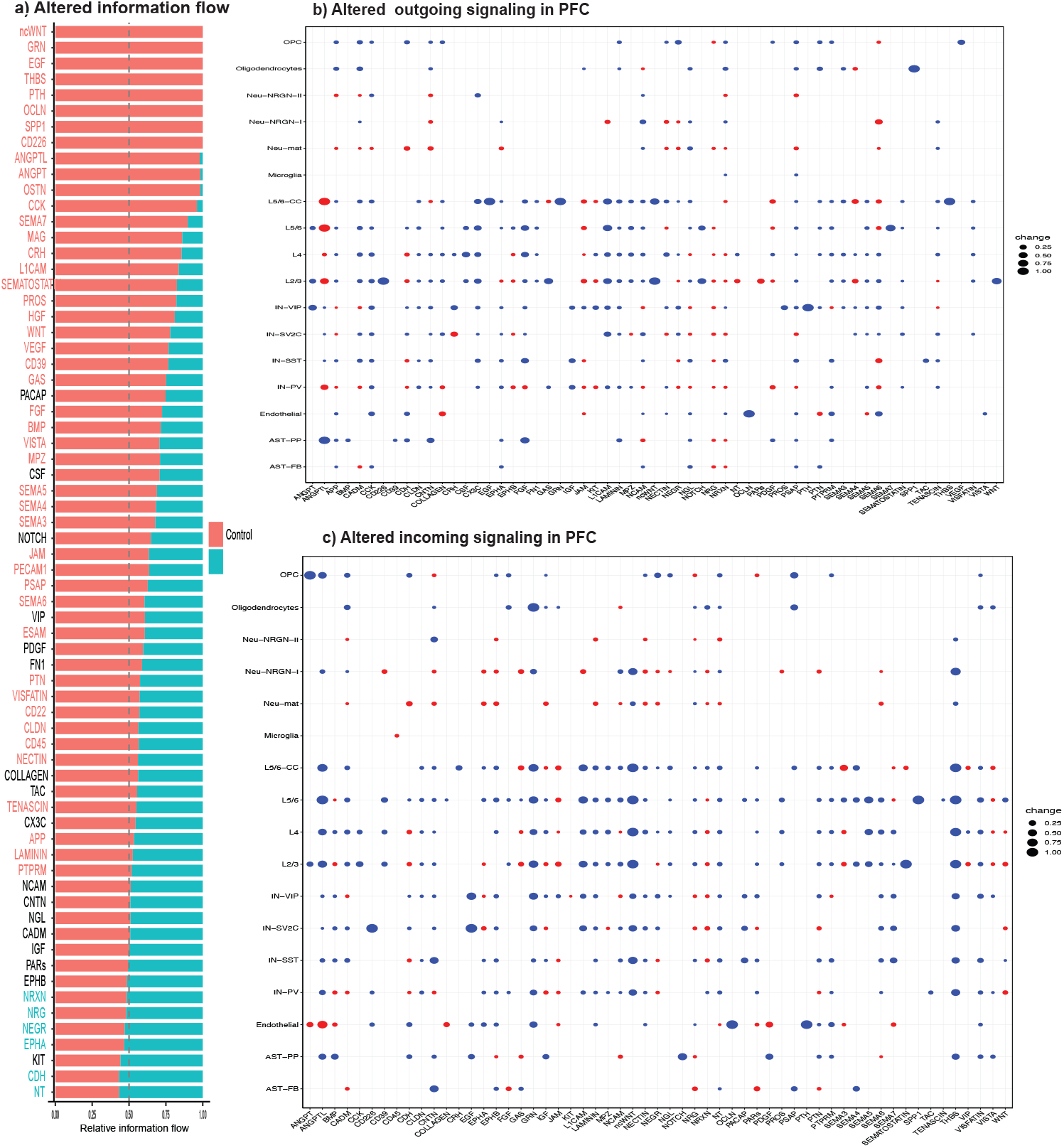
Comparison of pan cell type signaling networks in PFC. a) Pan cell type relative information flow showing signaling pathways identified in ASD PFC and controls. The pathways with greater information flow in ASD or controls were in cyan or red, respectively, with black indicating no significant differences. b,c) Dot plots showing the difference in relative contribution of each cell type to outgoing (b) or incoming (c) signaling in ASD vs controls. Signaling on the left (a) with no difference was not included in b or c.

In summary, our global analysis identified cell-cell communications and intercellular signaling that are potentially disrupted either between specific pairs of cell types or across many cell types in the ASD brains.

### Sample-by-sample CCC analysis in PFC

The above global approach is standard in CellChat and is also the most common design in cell-cell communication analysis by other software (i.e., only making two sample comparison). One concern is that it did not account for the potential differences among individual ASD or control samples. Furthermore, one can even argue that cells in one brain would never talk to cells in another. Thus, we decided to apply a complementary sample-by-sample approach. In essence, we used CellChat to obtain strengths of L-R interactions or score signaling pathways for each of the 13 ASD and 10 control PFC samples, as above, but then applied Wilcoxon test to determine if the sample-level strengths were significantly different between ASD and control groups. We focused our analysis on pan-cell type signaling pathways. In addition to the 68 identified by the above global approach, 24 others were detected. Of these 92 pathways, 10 were specific from controls and 3 unique to ASD. After pathways found in less than 5 samples were excluded, 74 were retained for ASD vs control comparison. A principal component analysis (PCA) of the samples based on the scores of the 74 signaling showed that the first and second components explained 44.7 % of the variation (**Figure S3)**, with noticeable but not completed separation between the ASD and control samples, indicating no major shifting of intercellular signaling flow between the two groups, consistent with results above. To gain better insight into the underlying molecular interactions, we decided to analyze all the L-R interactions in the 74 signaling by Wilcoxon test (p < 0.05). The results indicated that 88 L-R interactions in these signaling were significantly different between ASD and controls (**Figure 4**). These 88 L-R interactions were mapped to 32 signaling pathways, 25 and 7 of which were down and up regulated in the ASD PFCs, respectively, indicating that the other 42 signaling did not have significantly differential L-R interactions by our method. The data in **Figure 4** also help explain the apparent difference between an increase in pairwise cell-cell interaction strengths (**Figure 1c**) and a reduction of global CCC information flow in ASD (**Figure 3a**); there were more individual L-R interactions showing decreased strength (blue in **Figure 4**) but they involved fewer cell types than those interactions exhibiting increased strength (red in **Figure 4**).

**Figure 4.**
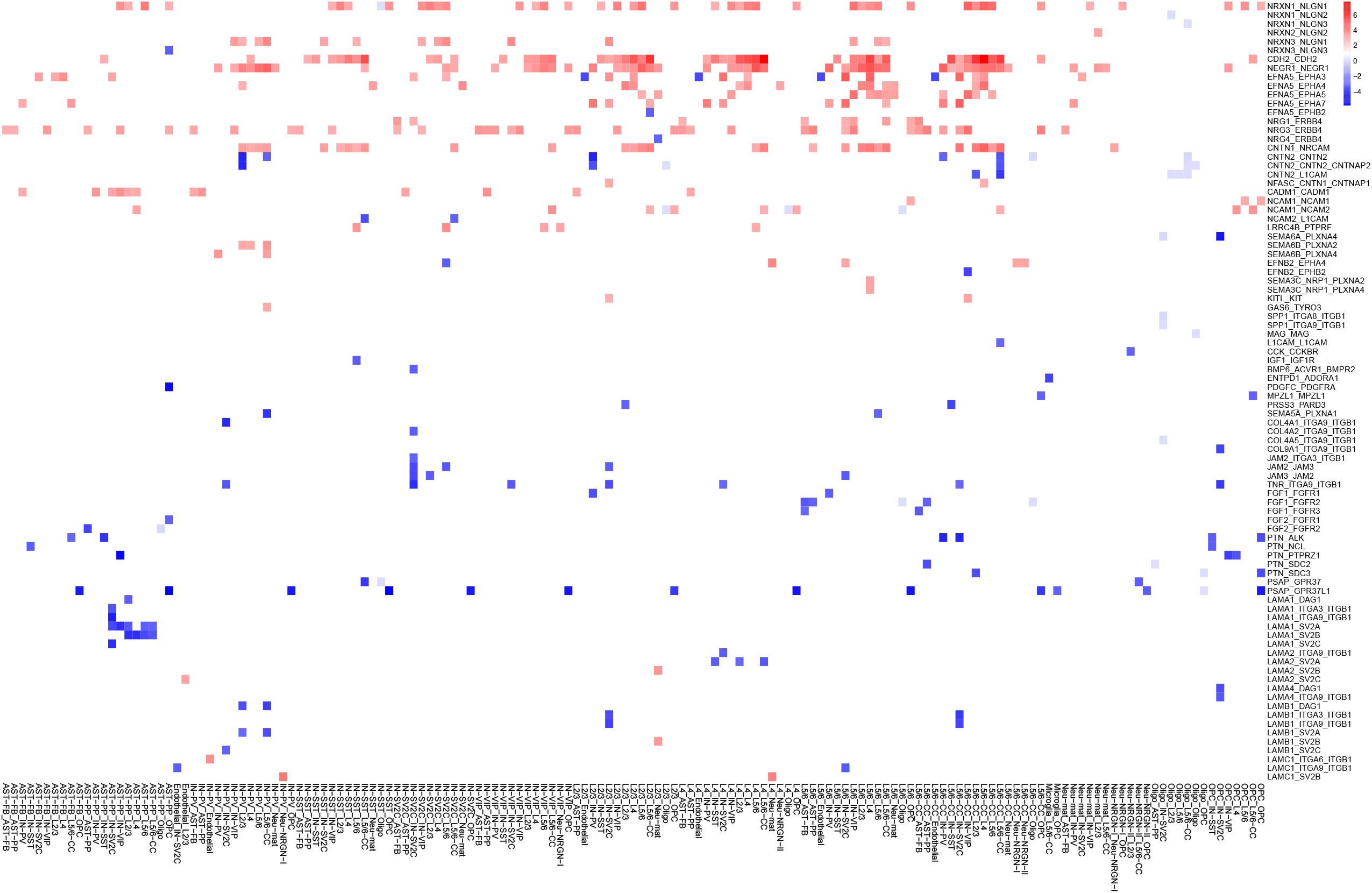
Heatmap for differential L-R interactions (y-axis) identified for individual pairs of cell types (x-axis) from the sample-by-sample approach.

From the 32 differentially active pathways, we selected seven with strong literature support for their potential involvements in ASD to illustrate our finding (see Discussion for details). They are neurexin (NRXN), fibroblast growth factor (FGF), contactin (CNTN), neural cell adhesion molecules (NCAM), netrin-G ligand (NGL), neuregulin (NRG), and pleiotrophin (PTN) signaling. Note that some of them (e.g., CNTN and NCAM) did not reach statistical significance in our global information flow analysis, indicating a difference in the statistic power of the two approaches. A chord diagram reflecting differences in interaction strengths between ASD and control samples are drawn for each pathway (**Figure 5**), including the directions of changes and the cell types involved. For each of the pathways, we also included the genes if their expression levels were determined to be significantly different between ASD and controls cells by Velmeshev *et al*.^23^. Among the seven, cell-cell interactions in NRXN and NRG were mostly increased in ASD, same for FGF and PTN, but similar numbers of increased and decreased interactions were observed in the other three pathways.

**Figure 5.**
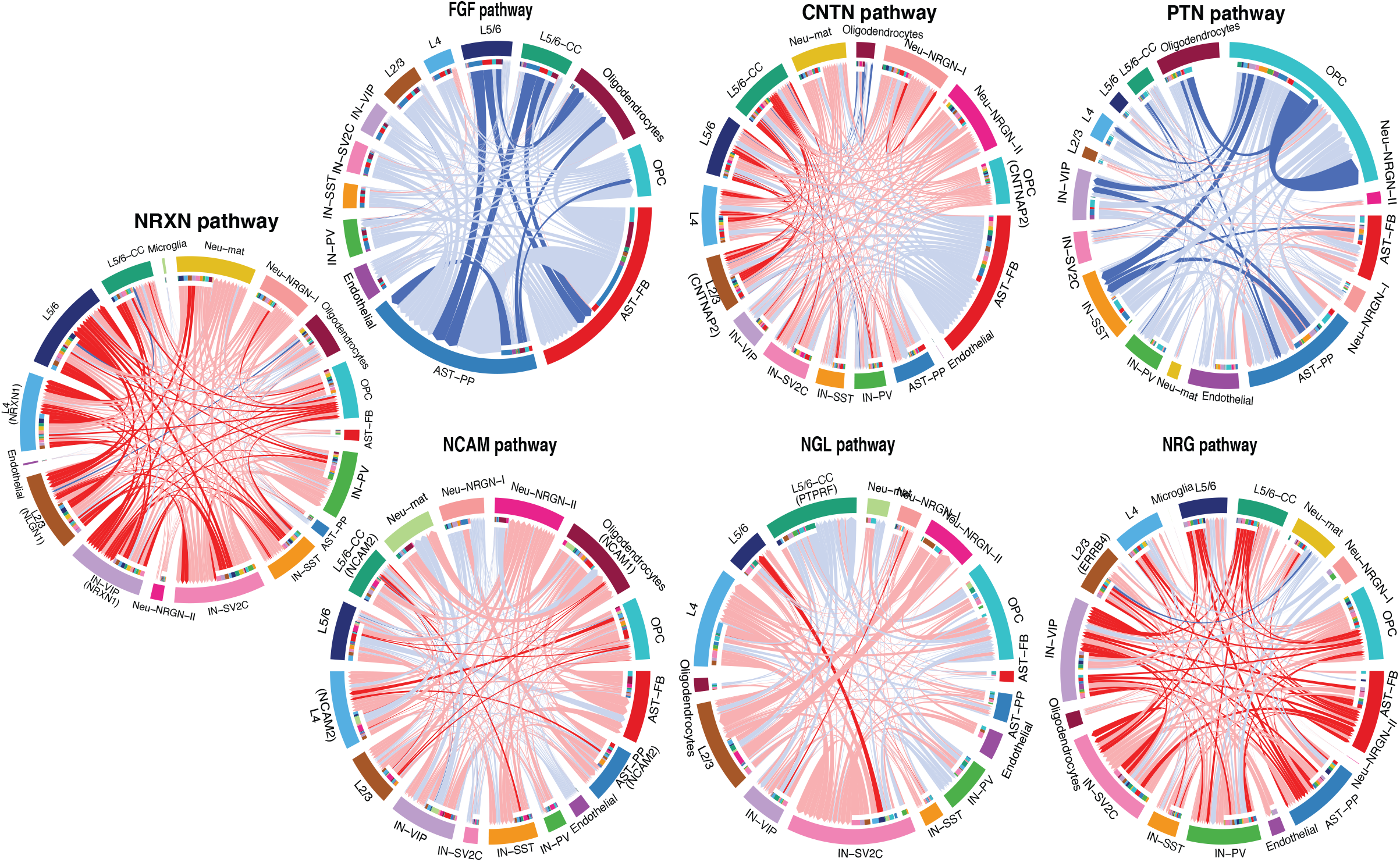
Chord diagrams plotting signaling strength differences between ASD and control PFC. The lines represent changes in L-R interaction strengths, with the statistically significantly different ones colored as intense red or blue, for increase or decrease in ASD, respectively. Light red or blue for small changes not reaching statistical significance. Grey lines for no changes. Genes identified as differentially expressed in Velmeshev *et al*.^23^ study were indicated in the corresponding cell type(s). The color bars in the inner circles indicates targeting cell types of the outgoing signaling while non-color part for incoming signaling.

In short, the results from our sample-by-sample approach strengthen our findings by the global method. Together, they highlight important intercellular signaling potentially disrupted in ASD PFC and demonstrate the advantage of CCC-focused analysis over conventional different gene expression analysis, as it can uncover altered intercellular signaling that would otherwise be missed in the latter.

### Function enrichment analysis of the disrupted cell-cell signaling

To better understand the roles of the disrupted cell-cell signaling, we carried out GO enrichment analysis, using 165 genes in the 32 dysregulated pathways. At FDR < 0.05, we found that the top “Biological Process”-related GO terms were axonogenesis, extracellular matrix organization, peptidyl-tyrosine phosphorylation, synapse organization, and regulation of cell morphogenesis, among others (**Figure 6a)** The enrichment analysis was reproduced by another software, ClueGO^35^, which also reduced redundancy in the enriched terms. As shown in **Figure 6b**, ClueGO was able to identify 8 clusters of GO terms, in which the main cluster, covering 27% of the terms, was related to axon development. This cluster is composed of neurogenesis, axon guidance, generation of neurons, neural crest migration or axon development terms, pathways with important roles in neurodevelopmental disorders. The second most enriched cluster (24% of terms) was “enzyme linked receptor protein signaling pathway,” in which terms such as MAPK cascade, ERK1 and ERK2 cascade or protein tyrosine kinase activity were included. Regulation of neuron projection development covered 8% of the terms, comprising key pathways related to developmental disorders such as regulation of neurogenesis, positive regulation of axonogenesis or regulation of nervous system development. Positive regulation of epithelial cell proliferation, biomineral tissue development, peptidyl-tyrosine phosphorylation and vasculature development accounted for 9%, 5%, 4% and 4% of terms, respectively. Taken together, the GO enrichment results are consistent with previous findings that have implicated these same cellular processes in ASD and other psychiatric conditions^10,11,13^, but put a more clear connection to the important roles of cell-cell communication.

**Figure 6.**
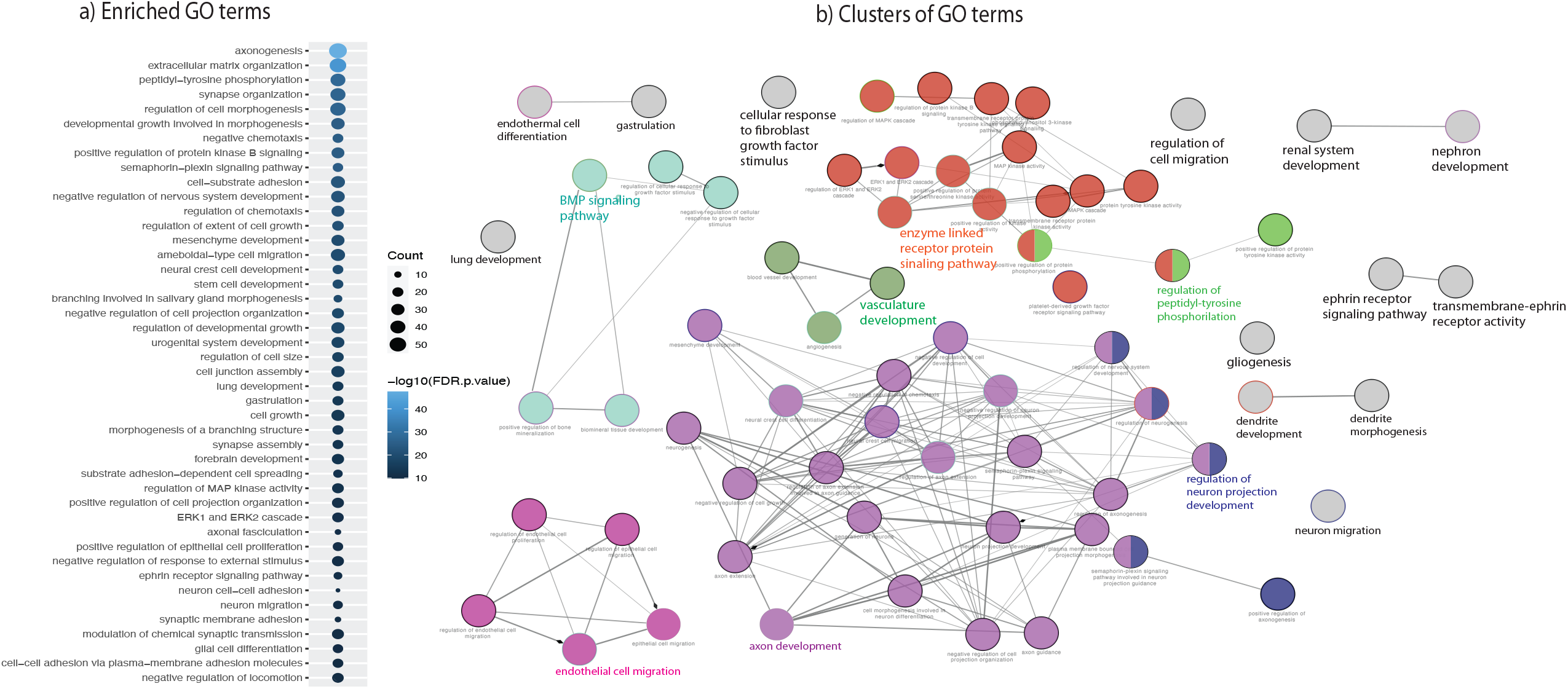
Function enrichment analysis of genes in the dysregulated CCC signaling. a) Dot plot showing the enriched GO terms. b) Network connecting GO terms with sharing genes. Nodes are enriched GO terms while the edges represent the extents of genes shared between two terms.

### Functional connection of the disrupted cell-cell signaling to ASD risk genes

In order to study how the disrupted cell-cell communication signaling are linked to ASD genetic risk, we applied the STRING database to construct a protein-protein interaction network. It identified 1,076 edges connecting 153 of the 165 genes in the disrupted CCC signaling (ranging from 1-30 connections) to 239 ASD risk genes in the SFARI database (**Figure 7a**). In addition, the network has an average local clustering coefficient of 0.38 and interaction enrichment values of 0.05, indicating overall strong connections among the genes (i.e., nodes).

**Figure 7.**
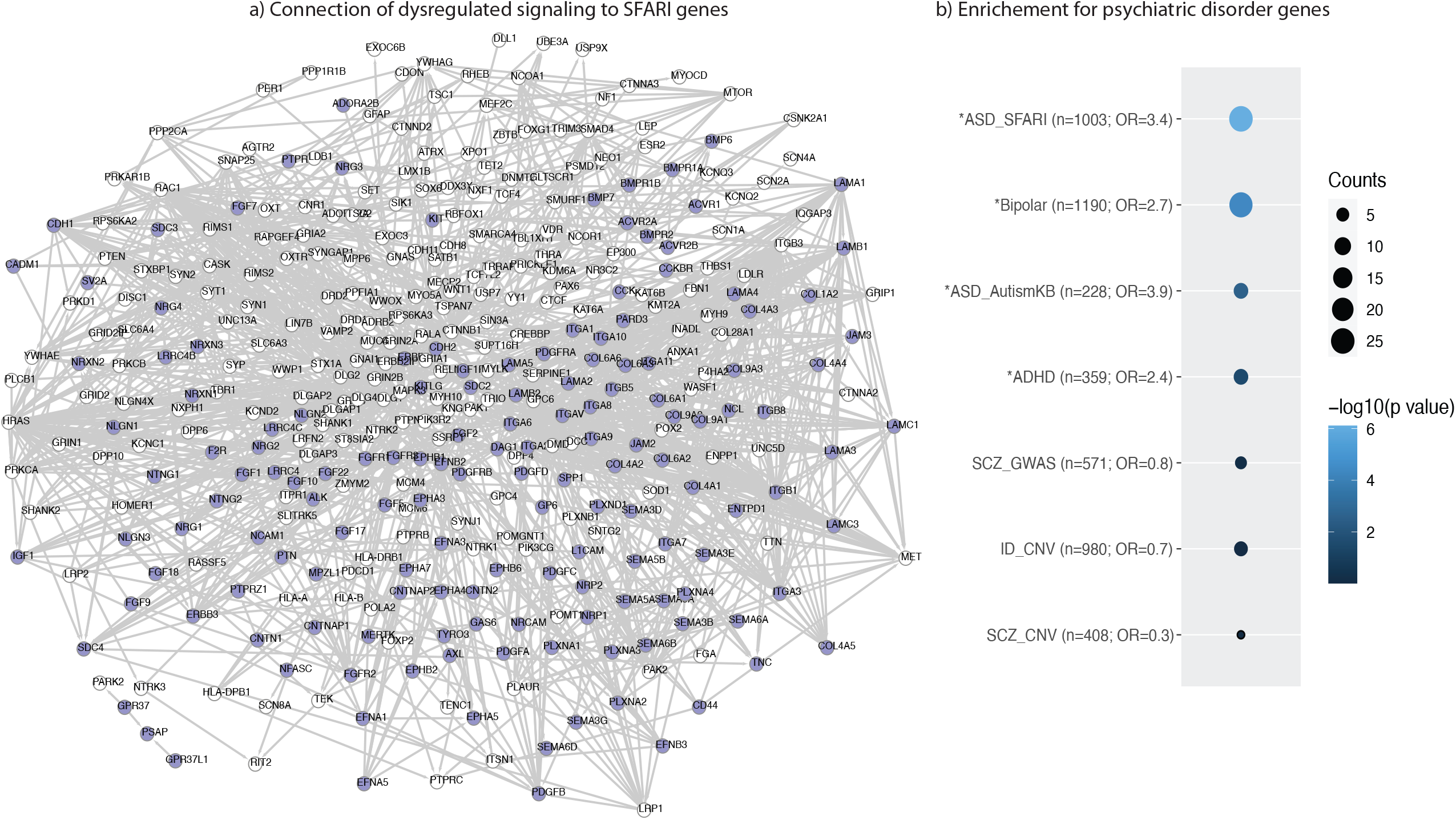
Connection between brain disorder risk genes and genes in the dysregulated CCC signaling. a) Protein interaction network connecting dysregulated CCC signaling genes (blue) and ASD risk genes (white) in SFARI database. b) Dot plot showing overlapping results of dysregulated signaling genes with lists of genes implicated in different brain disorders.

### Enrichment for gene sets related to brain disorders

To further address how disruption of the CCC signaling may be related to ASD, we intersected the 165 genes with lists of genes associated to ASD, schizophrenia (SCZ), intellectual disability (ID), bipolar disorder (BD), and attention deficit hyperactivity disorder (ADHD) and tested for significance of over representation (**Figure 7b**). Among the genes overlapping with ASD lists, *CNTNAP2, LRRC4C, NLGN2, NLGN3, NRXN1, NRXN2, NRXN3*, and *PLXNA4* are considered as ASD risk genes with “high confidence” in SFARI database, due to multiple studies linked them to Autism and other neurodevelopmental disorders^13,46–50^. Also, significant enrichment was found with bipolar disorder (odds ratio (OR)=2.7) and ADHD (OR=2.4) gene lists, similar to previous findings^12,51^. In addition to the ASD genes, *SEMA5A* or *NRG1* are also very interesting. Semaphorin 5A gene has been linked to Autism^52^ and additionally the failure of its expression has been linked to abnormal development of the axonal connections in the forebrain^53^. Decreased expression of neuregulin 1 has been linked to orchestration of a number of executive functions in ASD patients^54^, as well as SCZ and bipolar disorders^55^. Regarding ADHD enriched genes, Krumm *et al*.^56^ related 3 semaphorin receptor protein (PLXNA3) to ASD. This receptor is known to be important for axon pathfinding in the developing nervous system^57^. Even though there were overlaps of genes in the ID and SCZ lists, no statistical significance was found for the overlap between the 165 CCC genes and genes related to these two disorders.

### Relationship between disrupted intercellular signaling and differential gene expression programs

To address how disruption of the CCC signaling may lead to downstream gene expression difference in the ASD brains, we studied how they could be linked to the pathways enriched among DEGs in each of the cell types in the ASD brains. To do this, we first ranked genes by their expression fold changes between ASD and control cells in each of the 17 cell types and then performed GSEA. From the result, we select and show the enriched GO terms containing at least one gene in our disrupted CCC signaling **(Figure 8)**, resulting in 58 GO terms linked to 14 signaling pathways. 14 of the GO terms were more active in the controls, mostly involving non-neuronal cells. Oligodendrocytes showed 8 enriched pathways, including transmission of nerve impulse, regulation of glial cell differentiation, and cell chemotaxis. Similarly, AST-PP was enriched in integrin mediated signaling pathways and regulation of smooth muscle cell differentiation. GO terms more active in ASD samples were mainly seen in neuronal cells, i.e., IN-SST, IN-SV2C, IN-VIP, L2/3 and Neu-NRGN-I. In total, 45 enriched GO terms more active in ASD neurons were connected to disrupted CCC signaling, including synapse assembly, neuron cell adhesion, learning, regulation of neuron differentiation, postsynaptic specialization organization, receptor clustering, or synaptic signaling. Taken together, these results indicate that disrupted intercellular communications can be linked to downstream intracellular pathways and gene expression programs dysregulated in both neurons and glia cells in the ASD brains.

**Figure 8.**
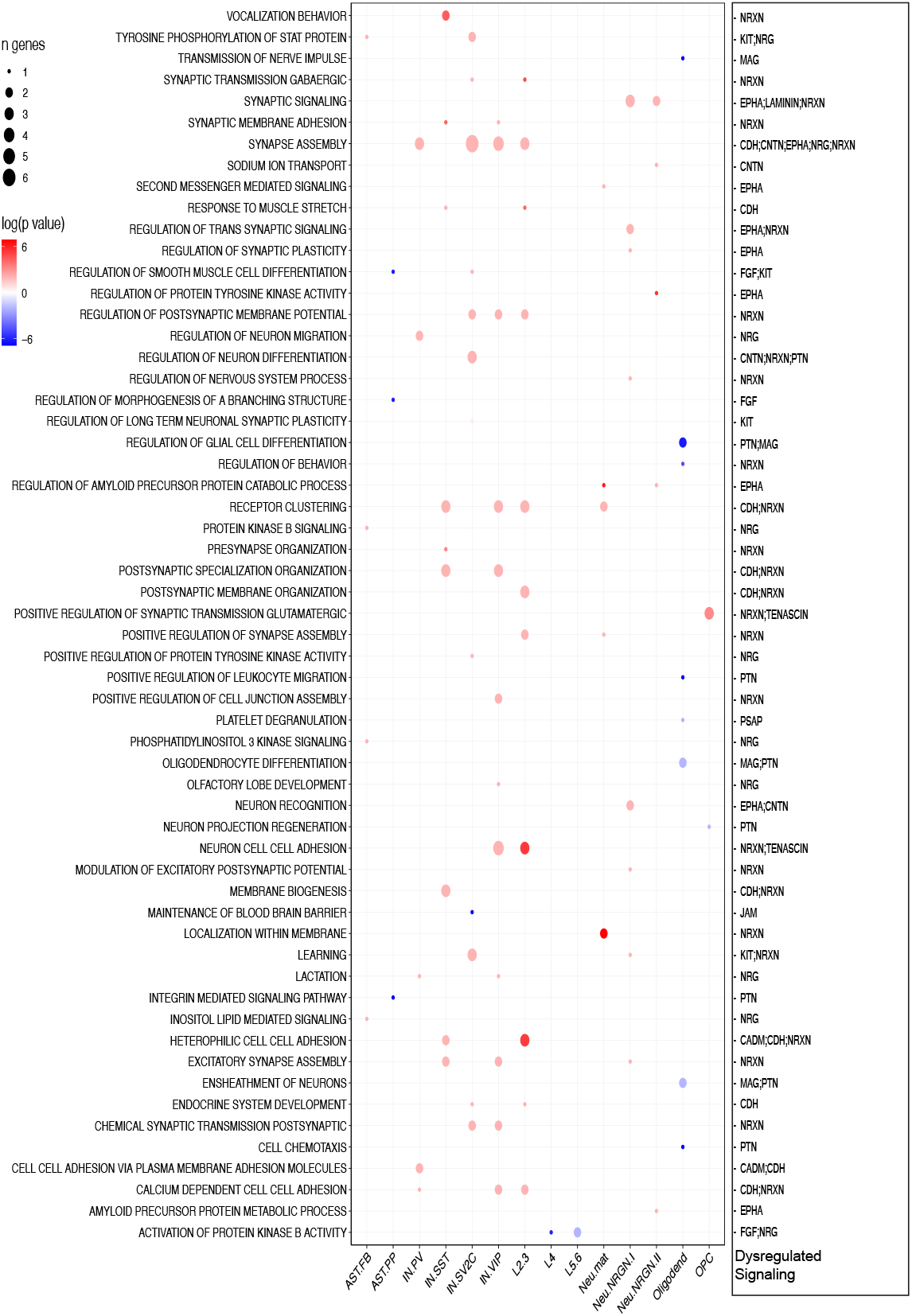
Connection between cell type ASD enriched pathways and dysregulated CCC signaling. Dot plot showing enriched pathways from GSEA for individual cell types, with red and blue for higher activities in ASD and control PFC, respectively. The right column lists the corresponding dysregulated signaling from CCC analysis.

### Spatial co-expression of ligands and receptors

Our analysis assumed that cells of the same type express the same ligands and receptors, by using snRNA-seq data that did not directly include spatial locations of cells, although the excitatory neurons did contain brain layer information. Spatial location, however, could be an important factor in cell-cell communication. To address this, we analyzed the spatial transcriptomics data from Maynard *et al*.^41^, which applied 10x Genomics Visium platform to generate a map of gene expression in the six-layered dorsolateral prefrontal cortex of an adult human brain. In their study, they have already corroborated their spatial registration to the cell type annotation in the snRNA-seq data used in our analysis. Here, we wanted to determine if the spatial expression levels of the ligand-receptor pairs are more correlated than expected by chance. The data indicate that most of the randomly paired L-R had correlation coefficients near “0” but the interacting pairs identified by CellChat showed significantly larger correlations, using spatial data across all brains or the subset in defined layers (**Figure 4Sa/b**). The correlation shifted to both positive and negative perhaps because CellChat interaction includes soluble agonists, antagonist, and co-stimulatory and co-inhibitory co-factors.

### Cell-cell communications in anterior cingulate cortex

We present our findings for PFC above, but we have also carried out analyses for the anterior cingulate cortex samples using our global approach. Similar to PFC, most of the cell-cell interactions in ACC also occurred through neuronal cell types (e.g., L5/6-CC and L2/3 excitatory neurons), while microglia and Neu-NRGN-I contributed the least. In contrast to PFC, our global approach showed a mixture of lost and gained pairwise interactions in the 17 ASD vs controls as mentioned above (**Figure 1d; Figure S5a**). L4, Nue-mat, Neu-NRGN-I, Neu-NRGN-II and AST-PP cell types showed an increased number of interactions, acting as receiver cell types, while the remaining cell types showed decreased interactions in ASD. Similar to PFC, CCC strengths **(Figure 1e; Figure S5b)** were increased in ASD ACC for most of the cell types. For pan cell type signaling network, CellChat identified 70 unique signaling pathways in ACC, with 5 specific to controls (ANGPT, CD226, CD99, ENHO and ncWNT) and 4 to ASD (EDN, FN1, PTH and TENASCIN). In **Figure S6**, we show the relative information flow for these pathways, with 37 significantly up- and 16 down-regulated in the ASD. Only 13 pathways showed a decreased activity, while 5 showed increased activity in both ASD PFC and ACC. Further analysis compared the relative contributions from each cell type for outgoing and incoming signaling in ASD and control ACC, indicating global similarity but noticeable changes for some pathways, such as CD99, ncWNT or ANGPTL (**Figures S7a/b**).

## Discussion

ASD is a neurodevelopmental disorder involving genetic, epigenetic, and environmental factors through various processes^3,58^. At the cellular level, cell-cell signaling is critical. Using CellChat, we have performed a CCC analysis using previously published single cell datasets^23^. The availability of data from both ASD and controls allowed us for the first time to systematically compare cell-cell communications in ASD and control brains. Results from our multiple-level CCC analyses suggest that several cell-cell signaling pathways are potentially disrupted in the ASD brains, involving mostly neurons, but glial cells also contribute. Our findings are in line with a large body of literature that support the importance of cell-cell interactions in neurodevelopment and brain disorders. For example, studies suggested that genes dysregulated in ASD were enriched in cell-cell communication, nervous system development, or cilia regulators or components terms^10,11^. Our computational analysis is based on the combined expression of ligands, receptors, and their cofactors in any pair of cell types. It is conceptually different from differential expression analysis based on individual genes. This can be seen in the results shown in **Figure 5**, where many of the disrupted signaling pathways contain only few or no DEGs that were determined from the same datasets.

Also different from many other cell-cell communication studies, we applied a combination of global approaches and sample-by-sample approaches. While the former potentially has higher sensitivity (due to more cells included), the latter likely has better precision as it requires independent statistical supports from many samples. Nevertheless, the disrupted cell-cell communications and the corresponding signaling pathways from the two approaches generally agree, with nearly all in the global approach also found by sample-by-sample approaches. For some signaling, the two approaches reported inconsistent results, probably because the degree of changes between ASD and controls is moderate, but further study will be needed. At the signaling level, our study indicates that more cell-cell communications show increased activities or strengths in ASD, but those involved in fewer cell types are decreased. Genes in these differential CCC pathways were enriched for molecular pathways and cellular processes that play important roles in neurogenesis and neuron functions, as well as risk genes for ASD.

For reproducibility, we performed a subsampling analysis by randomly selecting 50% of the cells (repeating 10 times). Analyzing the data by our global approach showed that the changes in L-R strengths between ASD and controls were highly correlated to what were obtained using all cells (**Figure S8**). In addition, we tested two other software (NATMI and CellPhoneDB)^29,31^ and obtained significantly overlapping but not identical results, with the difference largely due to differences in both the collection of ligand-receptors and how interactions are quantified, as discussed by others^33^.

To put our results in a broader literature content, below we discuss key signaling pathways with strong literature support for their association with ASD or abnormal neurodevelopment: NRXN, FGF, CNTN, NCAM, NGL, NRG and PTN pathways (**Figure 5**).

The neurexin pathway is composed of *NRXN1, NRXN2, NRXN3, NLGN1, NLGN2* and *NLGN3* genes, all of which except *NLGN1* are identified as “high confident” ASD genes in SFARI database. Many studies have also linked these genes closely to ASD and other neurodevelopmental disorders. Wiśniowiecka-Kowalnik *et al*.^59^ described three CNV within *NRXN1* in subjects from three families showing ASD, anxiety and depression, developmental delay, and speech delay. Furthermore, Pinto *et al*.^60^ detected an excess of exonic *NRNX1* CNVs in 996 cases compared to 4,964 controls. However, Wang *et al*.^47^ did not find an association between *NRXN1* and ASD, suggesting heterogeneity of the disorder, but their study associated *NRXN2* and *NRXN3* to ASD and specific alleles to ASD severity. On the receptor side, a nonsense variant in *NLGN2* was reported in a patient with severe anxiety, obsessive-compulsive behaviors, autism, short attention span, global develop-mental delays, hyperphagia, obesity, macrocephaly and dysmorphic features^48^. Similarly, two studies found that intronic *NLGN3* variants were potentially linked to ASD^61^ with a male bias^62^. Furthermore, overexpression of both *NRXN* and *NLGN* can lead to an alteration in Excitatory/Inhibitory ratio due to an increase in the number of synapses^63^.

In mammals there are 18 different FGF ligands and 4 FGF receptors (FGFRs), many of which regulate cell proliferation, cell division, and neurogenesis^64^. Here, we identified 12 that showed (5 after Wilcoxon test) a difference between ASD and controls, all of which had a decreased strength in ASD PFC (**Figure 5**). FGF1 and FGF2, FGFR1, FGFR2 and FGFR3 were involved. While FGFR1 and FGFR4 are mainly expressed in neurons, FGFR2 and FGFR3 are expressed in astrocytes and oligodendrocytes^65^. In accordance with that, the only significant FGF1-FGFR1 interaction was found between L5/6 and IN-PV neurons, while the interactions with FGFR2 and FGFR3 receptors were found with astrocytes and oligodendrocytes as receiver cells. The FGF system is known to be involved in different brain-related disorders. Esnafoglu and Ayyıldız (2017), for example, reported significantly lower levels of FGF2 in ASD children. Additionally, Evans *et al*.^67^ linked decreased expression of FGF1, FGF2, FGFR2 and FGFR3 in cortical areas to depressive disorder, while Liu *et al*.^68^ and Wang *et al*.^69^ linked FGF2 and FGFR2 respectively to bipolar disorder.

The contactin pathway (CNTN), on the other hand, showed significant L-R interactions between most pairs of the cell types. Genes in these significant interactions were *CNTN1, CNTN2, CNTNAP1, CNTNAP2, NRCAM, L1CAM* and *NFASC*. Conactin-1 and Contactin-2 are important for neuron-glia interactions, and they regulate neuronal migration and axon guidance^70^. However, no direct link has been found to ASD^71^. Its interaction partners, *CNTNAP2* and *NRCAM*, on the other hand, were identified as syndromic and strong candidate and as gene with suggestive evidence, respectively. *CNTNAP2* is a very well-known gene in ASD, and brains with the risk variants show more connectivity between the frontal cortexes, while connections dimmish between more distant regions^72^. Also, Li *et al*.^73^ suggested susceptibility of the gene in the Chinese Han population, while more recently Nascimento *et al*.^74^ linked a single nucleotide polymorphism (SNP) of *CNTNAP2* in a Brazilian population. *Cntnap2* knockout rats exhibited deficits in sociability and social novelty, as well as, exaggerated acoustic startle responses, increased avoidance to sounds, and a lack of rapid audiovisual temporal recalibration ^75^, all of which are related to the symptoms in ASD.

Neural cell adhesion molecules (NCAM) play a key role in neural development, and shaping axonal and dendritic arborizations^76^. Its downstream signaling determines if a cell migrates or projects axons and dendrites to targets in the brain^77^. These molecules participate in NCAM-NCAM homophilic interactions as well as heterophilic interactions with molecules such as L1^78^ or FGFR^79^ among others. Even though FGFR receptors were also identified in this study, only three interactions, NCAM1-NCAM1, NCAM1-NCAM2 and NCAM2-L1CAM were determined as significantly different between ASD and controls. While some studies have linked decreased levels of NCAM1 to ASD^80,81^, Gomez-Fernandez *et al*.^82^ and Purcell *et al*.^83^ could not detect differences in plasma cytokines, adhesion molecules or growth factors in children with ASD, compared with typically-developing children.

The interaction between leucine-rich repeated-containing (LRRC) protein 4B, also known as NGL-3, and receptor-type tyrosine-protein phosphatase (FPTPRF/LAR) was the only one showing a significant change in the Netrin-G ligand (NGL) pathway in ASD PFC. NGL are synaptic adhesion molecules that regulate synapse development, function and plasticity, with three family members: NGL-1/LRRC4C, NGL-2/LRRC4, and NGL-3/LRRC4B^84^. Lee *et al*.^85^ suggested that NGL-3 regulates brain development, and locomotive and cognitive behaviors among others. However, no data suggests a link between *NGL-3* gene and ASD. The LAR complex, on the other hand, is part of the RPTPs large family known to be involved in broader functions in the central nervous system^86^. Even though, other RPTP complexes have been associated with ASD^60^, no direct information exists for LAR complex.

Neuregulin (NRG) complexes, NRG1/2/3/4, play important roles in many neurological disorders such as brain trauma, spinal cord injury, and SCZ^87,88^. All isoforms contain a fragment encoding epidermal growth factor (EGF)- like domain that enables the interaction with ErbB receptor tyrosine kinases^89^. Many studies have been published about these interactions regarding important functions in cell differentiation, neuronal migration, or synapse formation^90,91^. Three interactions between these complexes were among our significant results: ligands NRG1, NRG3 and NRG4, and receptor ErbB4. Many studies link the NRG1 complex to ASD. Abbasy *et al*.^54^, for example, reported that down regulation of the complex is linked to deficits in response, vigilance, and working memory. Likewise, Esnafoglu^92^ suggested that NRG1-ErbB signal system may contribute to the disorder, and Dabbah-Assadi *et al*.^93^ suggested that NRG1-ErbB4 pathway disruption can lead to cognitive dysfunction. While NRG2 and NRG3 complexes have not been linked to ASD, many reports support their link to other neurodevelopmental disorders such as SCZ and bipolar disorder^94,95^.

In the PTN pathways, the unique ligand from our analysis was pleiotrophin protein, an extracellular matrix-associated protein known to be involved in neurodevelopmental processes such as cellular proliferation, early presynaptic or postsynaptic specialization^96^. A previous study^97^ demonstrated that PTN knockout mice showed disruptions in cognitive and affective processes. Another study suggested that both PTN and its receptor, protein tyrosine phosphatase receptor type Z1 (PTPRZ1), could be potential candidate genes for ASD ^98^.

In addition to the above signaling pathways with literature support for their association to ASD, our unbiased over representation analysis also found GO “Biological Processes” terms related to axonogenesis, extracellular matrix organization, peptidyl-tyrosine phosphorylation, synapse organization or regulation of cell morphogenesis. Similarly, Gai *et al*.^99^ in their study of ASD risk genes from 631 autism subjects, 1,162 parents, and 1,775 control children found enrichments in synaptic functions, such as, cell-cell signaling, transmission of nerve impulse or neurotransmitter transport. Similar results were reported in an independent study^100^. In this study, 112 ASD genes were selected for enrichment analysis in mouse and human phenotypes. The results linked these genes to changes in brain and neuronal morphology, electrophysiological changes, neurological changes, and higher-order behavioral changes. In a comparative study between ASD and SCZ^101^, gene sets enriched in protein phosphorylation/kinase activity, among others, were found for ASD. Furthermore, this study suggested an etiological overlap of ASD and SCZ since 8% of the clinically significant CNVs were shared between the two disorders. This comes as no surprise since a large epidemiological study^102^ has found that a family history of SCZ is a risk factor for ASD. Moreover, in 50% of ASD cases there is an association with intellectual disability as well as comorbidity with other psychiatric disorders^58,103^. In this regard, it is interesting that our analysis of the disrupted cell-cell communication genes found no enrichment with SCZ and ID genes but positive enrichments for bipolar disorder, ADHD and ASD.

In this study, cell-cell communication is inferred from the expression of protein-coding genes thus not capturing other signaling events in the brain, such as non-protein molecules (e.g., neurotransmitters). Another limitation of the study, as in other cell-cell interaction studies based on scRNA-seq or snRNA-seq, lies in the lack of spatial information, as juxtracrine and paracrine signaling operate from 0 to 200 μm^104^. The application of spatial transcriptomic data will help, but that technology by itself is currently not able to provide single cell resolution data. However, one study applied spatial transcriptomics to investigate alterations occurring in tissue domains within a 100-μm diameter around amyloid plaques in Alzheimer’s disease and found alterations consistent with disease pathogenesis^105^. As the technology advances and more data become available for ASD, we should be able to improve the methods in this study, as discussed in a recent review^104^. In addition, since the snRNA-seq data were from postmortem brains, our findings by themselves cannot distinguish if the CCC signaling disruption contributes directly to ASD pathogenesis or is a result of the brain’s response to ASD conditions. Lastly, signaling in the brain is conducted by proteins, while our analysis was based on RNAs.

## Conclusions

In this study we perform a comprehensive bioinformatics analysis to characterize the cell-cell communication changes in the brains of autistic subjects. We found multiple intercellular signaling pathways that are potentially altered in autism. Genes in these altered signaling pathways show a significant enrichment for genes involved in both ASD risk and with intracellular molecular pathways that are putatively dysregulated in multiple brain cell types.

## Supplementary Figure Legend

**Figure S1. Scatter plot for cell numbers in the 17 cell types and their numbers of CCC interactions**.

**Figure S2. Heatmaps showing CCC differences between ASD and control PFCs in terms of counts (a) and strengths (b)**. The rows and columns indicate signaling sending and receiving cells, respectively.

**Figure S3. PCA plot based on pathway interaction strengths in ASD and control samples**.

**Figure S4. Density plots for spatial expression correlation of actual L-R pairs identified in the scRNA-seq data or randomly paired L-Rs using either all spatial spots (a) or only spots in defined brain layers (b)**.

**Figure S5. Heatmaps showing CCC differences between ASD and control ACCs in terms of counts (a) and strengths (b)**.

**Figure S6. Pan cell type signaling networks identified in ASD ACC vs controls**.

**Figure S7. Dot plots showing the change in relative contributions of each ACC cell type to outgoing (a) or incoming (b) signaling in ASD vs controls**.

**Figure S8. Bubble plots showing the change in interaction strengths between individual pairs of cell types**. The y-axis shows the differences computed with all cells, while the x-axis shows the differences computed with one half of the cells. In the latter analysis, 50% of the cells were randomly selected to run CellChat 10 times, the mean differences are plotted on the x-axis with the standard deviations from the 10 runs indicated by bubble sizes.

## Ethics approval and consent to participate

Not applicable

## Consent for publication

Not applicable

## Competing Availability of data and materials

No new experimental data were acquired in this study and the snRNA-seq data are publicly available at (https://cells.ucsc.edu/?ds=autism).

## Authors’ Contributions

MA performed all the bioinformatics analysis and prepared the manuscript. HML contributed to analysis and edited the manuscript. DZ conceived of the experiment, contributed to the analytic plan, edited the manuscript, and supervised the work. All authors read, edited, and approved the final manuscript.

## Acknowledgements

We would like to thank members of the Zheng lab for valuable discussion, suggestions, and review of the manuscript.

