## supplemental figures for "Characterization of Cell-cell Communication in Autistic Brains with Single Cell Transcriptomes"

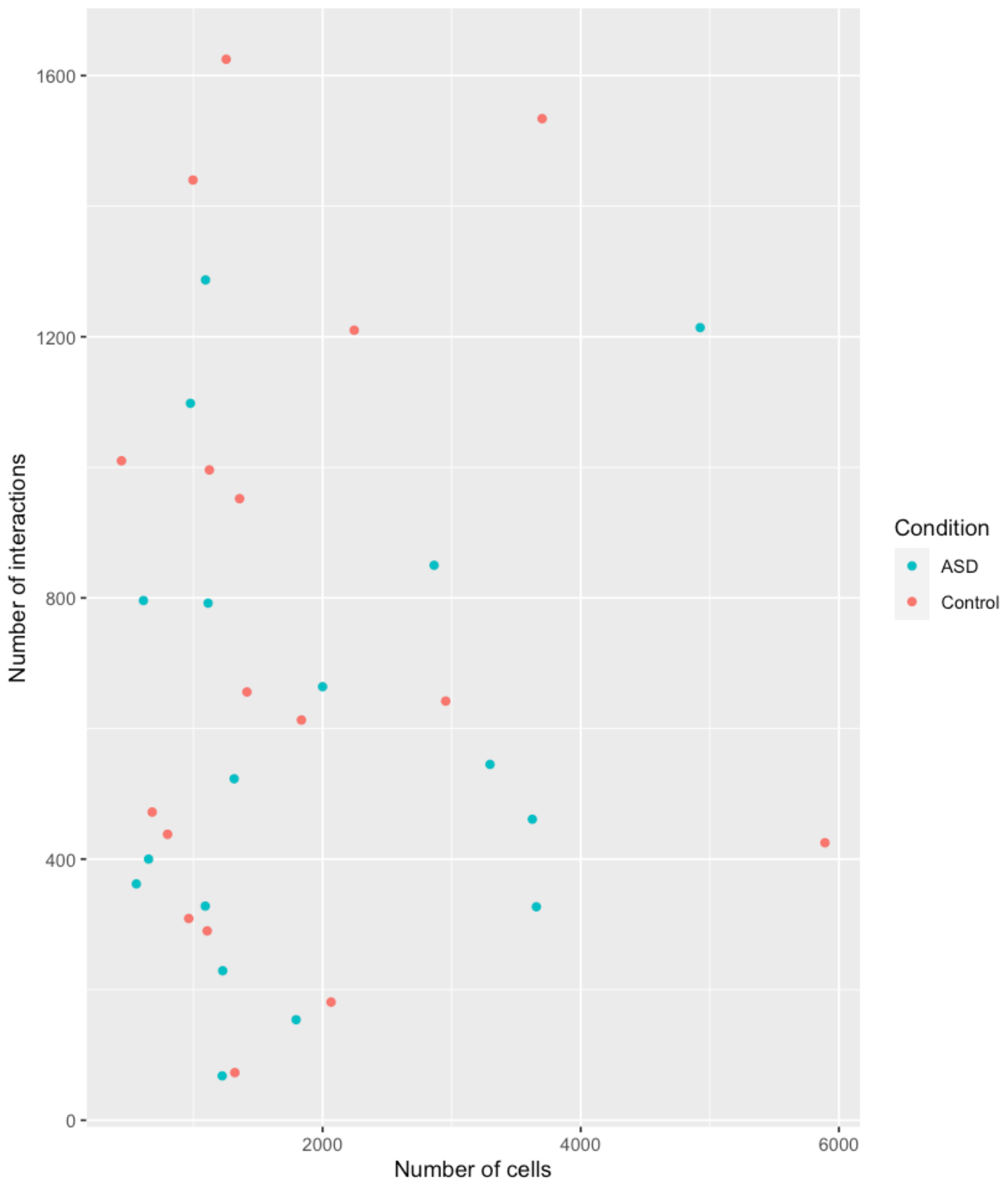

**Figure S1. Scatter plot for cell numbers in the 17 cell types and their numbers of CCC interactions.**

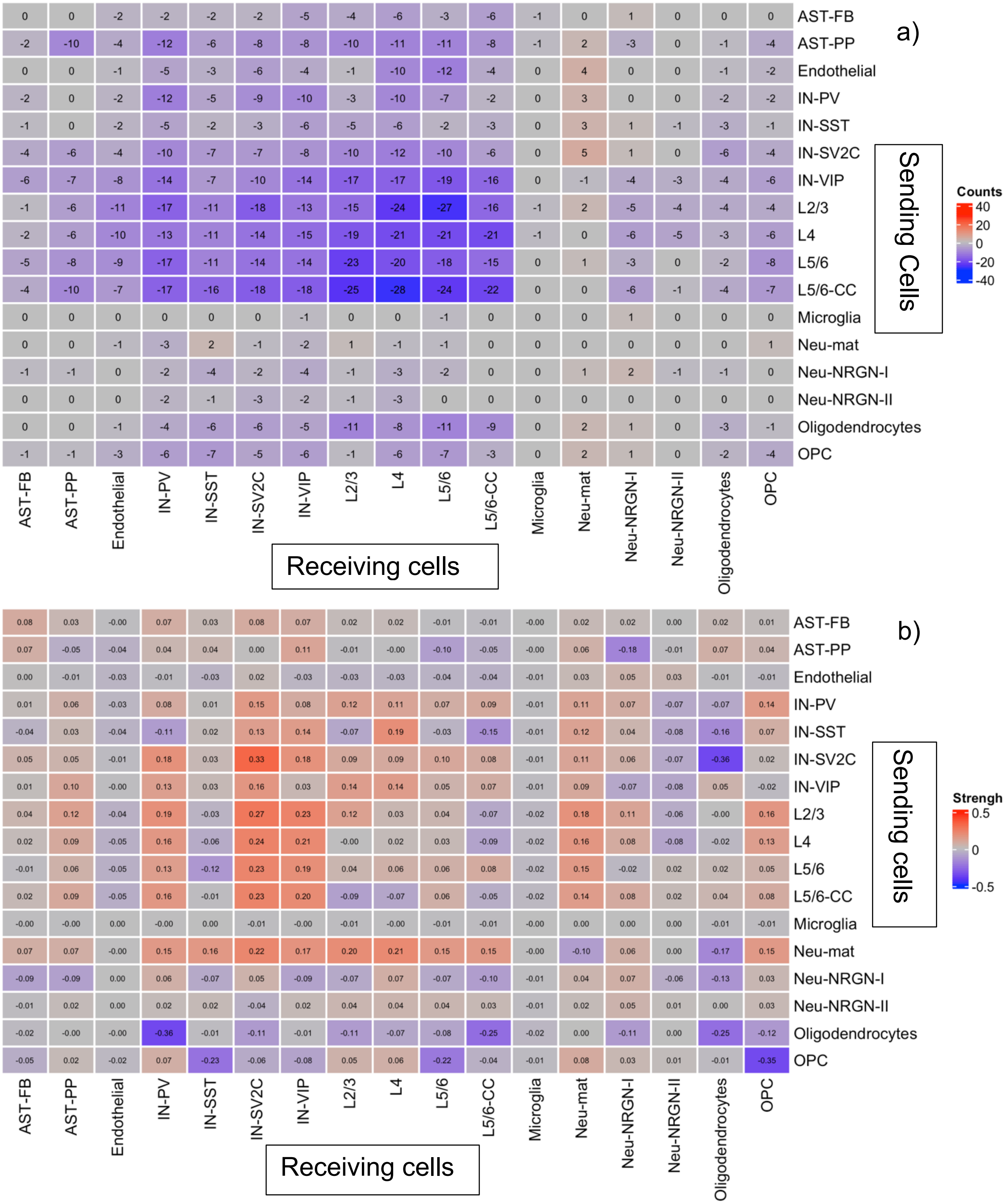


**Figure S2. Heatmaps showing CCC differences between ASD and control PFCs in terms of counts (a) and strengths (b).** The rows and columns indicate signaling sending and receiving cells, respectively.

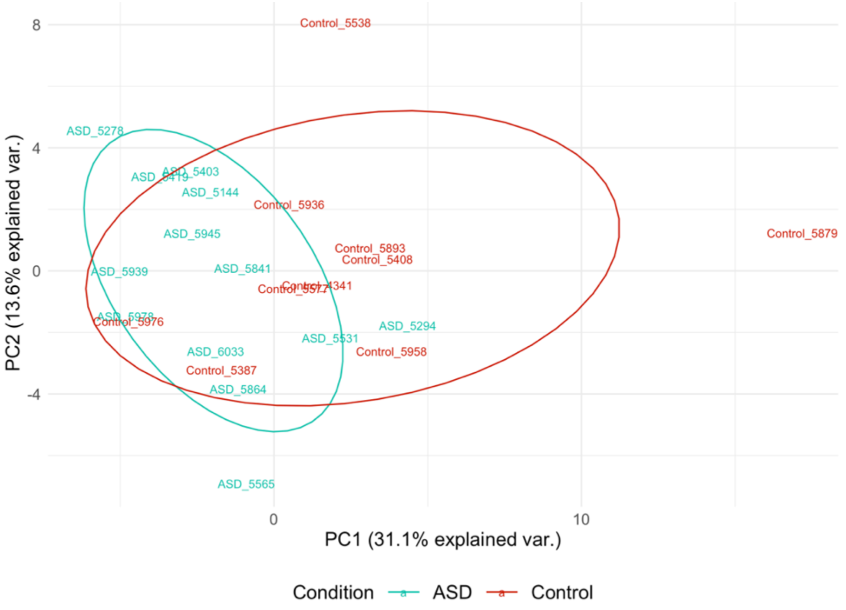


**Figure S3. PCA plot based on pathway interaction strengths in ASD and control samples.**

a)

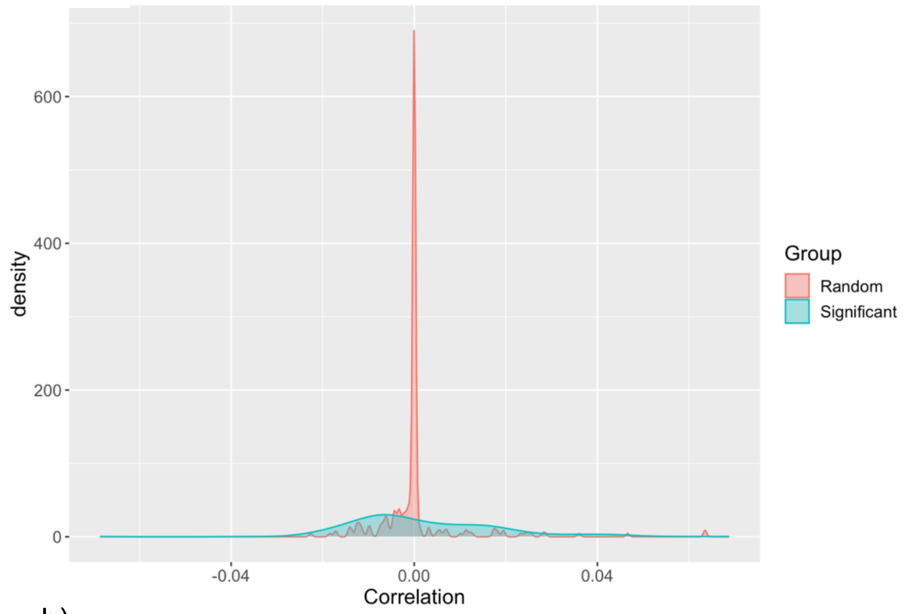


b)

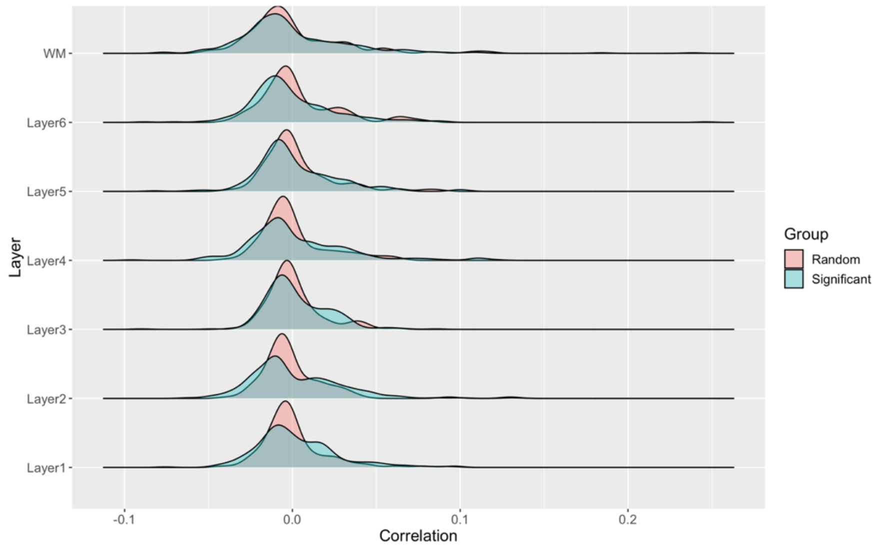


**Figure S4. Density plots for spatial expression correlation of actual L-R pairs identified in the snRNA-seq data or randomly paired L-Rs, using either all spatial spots (a) or only spots in defined brain layers (b).**

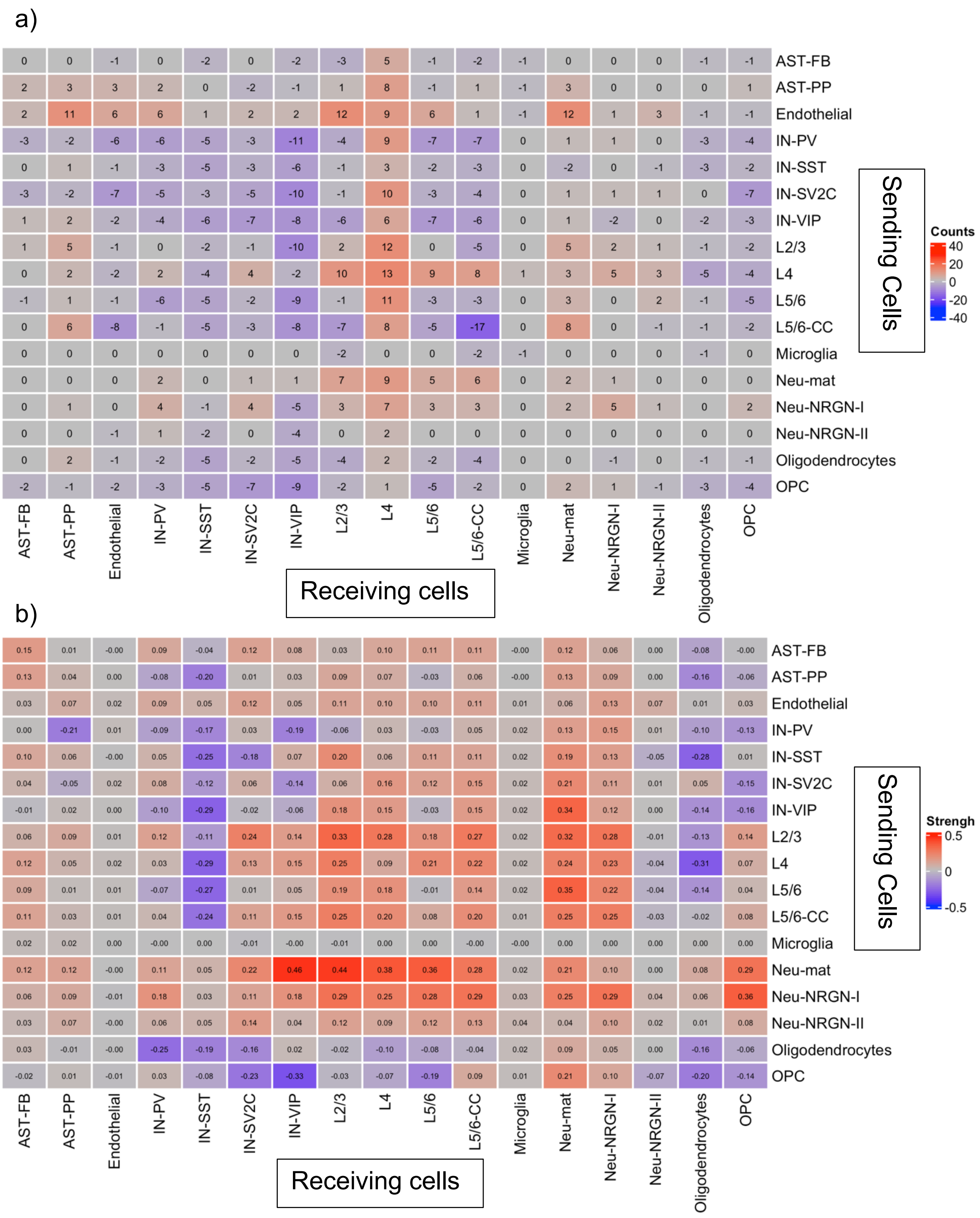


**Figure S5. Heatmaps showing CCC differences between ASD and control ACCs in terms of counts (a) and strengths (b).**

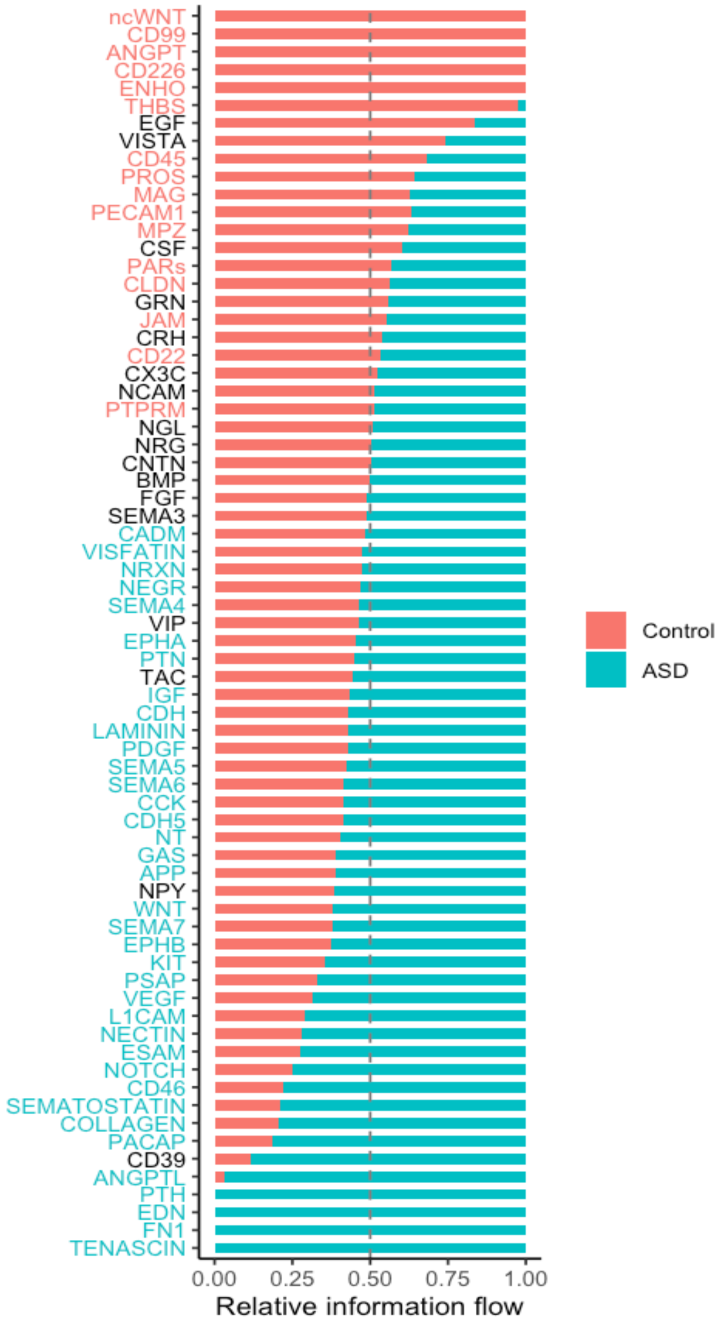


**Figure S6. Pan cell-type signaling networks identified in ASD ACC vs controls.**

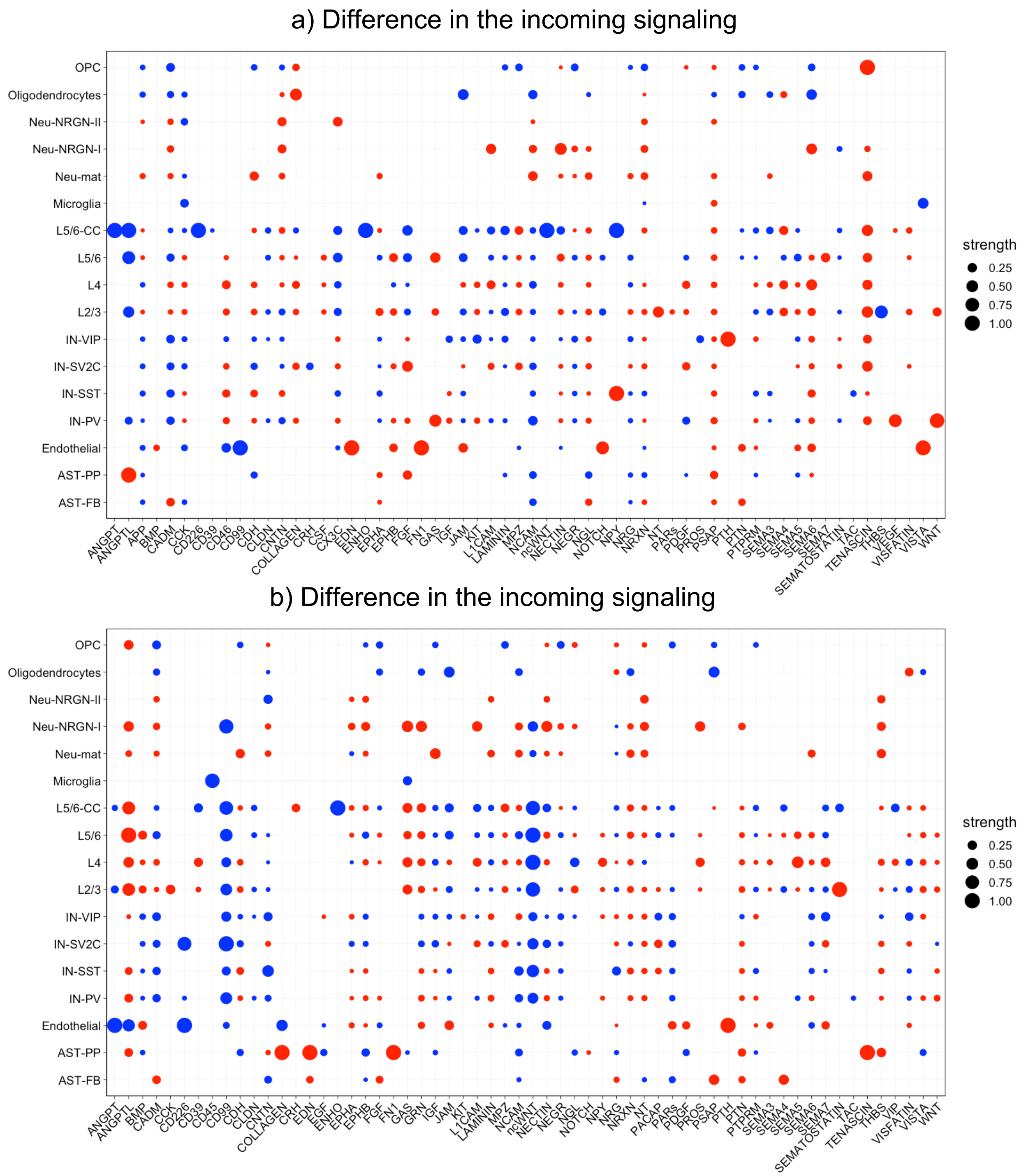


**Figure S7. Dot plots showing the change in relative contributions of each ACC cell type to outgoing (a) or incoming (b) signaling in ASD vs controls.**

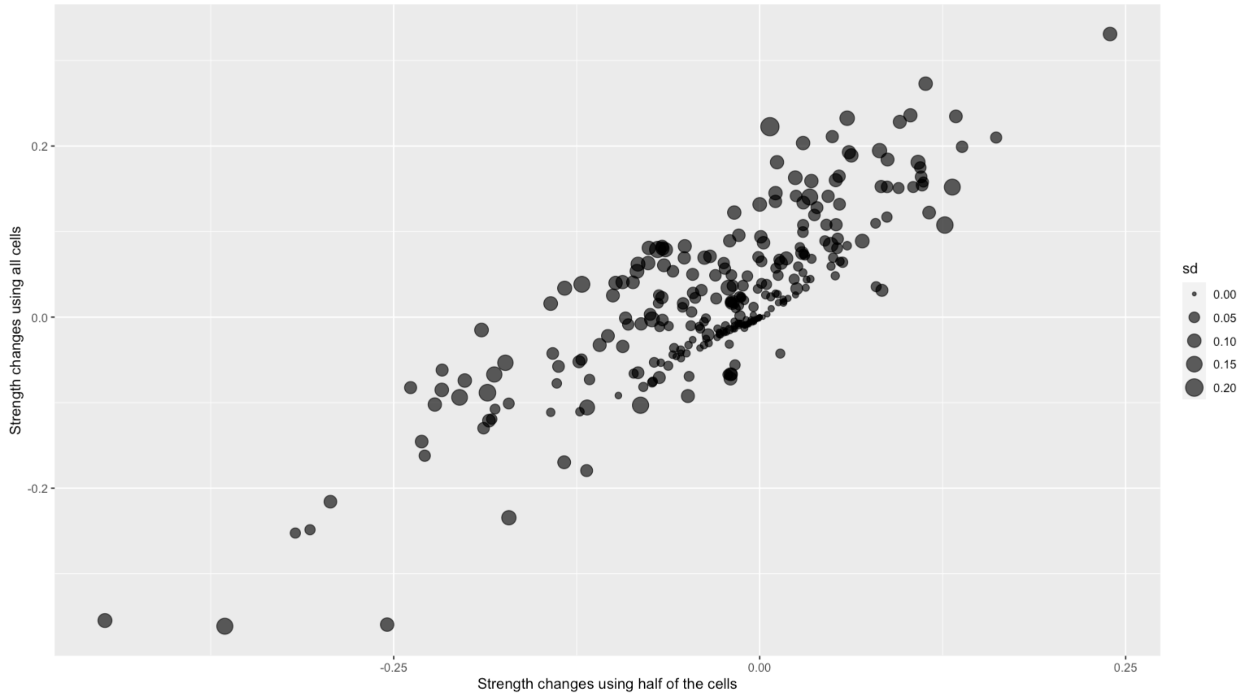


**Figure S8. Bubble plots showing the change in interaction strengths between individual pairs of cell types.** The y-axis shows the differences computed with all cells, while the x-axis shows the differences computed with one half of the cells. In the analysis of one half of the cells, 50% of the cells were randomly selected to run CellChat 10 times, the mean differences are plotted on the x-axis with the standard deviations from the 10 runs indicated by bubble sizes.
